# Long-range inhibitory intersection of a retrosplenial thalamocortical circuit by apical tuft-targeting CA1 neurons

**DOI:** 10.1101/427179

**Authors:** Naoki Yamawaki, Xiaojian Li, Laurie Lambot, Lynn Y. Ren, Jelena Radulovic, Gordon M. G. Shepherd

## Abstract

Dorsal hippocampus, retrosplenial cortex (RSC), and anterior thalamic nuclei (ATN) interact to mediate diverse cognitive functions, but the cellular basis for these interactions is unclear. We hypothesized a long-range circuit converging in layer 1 (L1) of RSC, based on the pathway anatomy of GABAergic CA1 retrosplenial-projecting (CA1-RP) neurons and thalamo-restrosplenial projections from ATN. We find that CA1→RSC projections stem from GABAergic neurons with a distinct morphology, electrophysiology, and molecular profile, likely corresponding to recently described Ntng1-expressing hippocampal interneurons. CA1-RP neurons monosynaptically inhibit L5 pyramidal neurons, principal outputs of RSC, via potent GABAergic synapses onto apical tuft dendrites in L1. These inhibitory inputs align precisely with L1-targeting thalamocortical excitatory inputs from ATN, particularly the anteroventral nucleus, forming a convergent circuit whereby CA1 inhibition can intercept ATN excitation to co-regulate RSC activity. Excitatory axons from subiculum, in contrast, innervate proximal dendrites in deeper layers. Short-term synaptic plasticity differs at each connection. Chemogenetically abrogating inhibitory CA1→RSC or excitatory ATN→RSC connections oppositely affects the encoding of contextual fear memory. Collectively, our findings identify multiple cellular mechanisms underlying hippocampo-thalamo-retrosplenial interactions, establishing CA1 RSC-projecting neurons as a distinct class with long-range axons that target apical tuft dendrites, and delineating an unusual cortical circuit in the RSC specialized for integrating long-range inhibition and thalamocortical excitation.

## Introduction

GABAergic projection neurons – inhibitory neurons with long-range axons – are important circuit elements in many brain areas, such the corticonuclear projections of Purkinje cells in the cerebellum and the striatonigral and striatopallidal projections of spiny projection neurons in the basal ganglia. In neocortical circuits, however, inhibition is mostly locally sourced, engaged through potent feedforward excitation of interneurons by excitatory afferents ^1^ Several exceptions have been identified, including inhibitory afferents to neocortex from somatostatin-expressing neurons in the zona incerta that project to sensorimotor cortex ^2, 3^, GABAergic neurons in the globus pallidus that project to frontal cortex ^4^, and parvalbumin-expressing neurons that project via corpus callosum to contralateral cortex ^5, 6^. For each of these, progress has been made toward characterizing both the presynaptic neurons and the long-range inhibitory circuits they form with postsynaptic cortical neurons.

A GABAergic projection from hippocampus to parietal cortex has also been described anatomically, arising from neurons in dorsal CA1 at the stratum radiatum/lacunosum-moleculare (SR-SLM) border that send axons to the retrosplenial cortex (RSC), ramifying in L1 ^7, 8^. Unlike the neocortically projecting GABAergic neurons mentioned above, however, the cellular properties of the presynaptic neurons are obscure, and the targets of their axons in the RSC are unknown.

This GABAergic CA1→RSC projection lies within a larger network thought to support the encoding and storage of hippocampally derived information in the RSC (reviewed in ^9-11^). Understanding how the GABAergic neurons in CA1 are connected to postsynaptic RSC neurons could thus yield mechanistic insight into the cellular basis for behaviors mediated by hippocampo-retrosplenial communication in this system.

Hippocampo-retrosplenial communication also involves excitatory circuits, but unlike GABAergic CA1→RSC projections these appear to be mostly indirect, through polysynaptic chains of connections ^12^. Two major routes carrying hippocampus-related information to the RSC are a corticocortical pathway via subiculum, and a subcortical pathway traversing the anterior thalamic nuclei (ATN). Subiculum→RSC projections arise from burst-firing vGlut1- or vGlut2-expressing subicular pyramidal neurons, which directly excite RSC pyramidal neurons ^13^. They also drive feedforward (disynaptic) inhibition ^13^, a local-circuit inhibitory mechanism previously implicated in mnemonic functions of the RSC ^10^. The ATN→RSC projections arise particularly from the anteroventral (AV) nucleus, and ramify mainly in L1 of the RSC ^14, 15^, similar to the GABAergic CA1→RSC projection. The cellular targets and circuits of ATN→RSC axons in the RSC are unknown, but the anatomical convergence in L1 of these inhibitory CA1 axons, combined with the presence of apical tuft dendrites of L5 pyramidal (L5pyr) neurons in this layer, suggests the possibility of a highly specific circuit configuration whereby long-range inhibition and TC excitation converge to co-regulate RSC output.

Here, we applied cell-type-specific optogenetic, chemogenetic, electrophysiological, and behavioral tools to address whether retrosplenially projecting CA1 neurons constitute a distinct class of long-range-projecting GABAergic neurons, and to test the hypothesis that they form CA1→RSC circuits that mediate direct inhibitory hippocampo-retrosplenial interactions, and converge with excitatory ATN-TC inputs via L1 circuit connections to modulate thalamo-retrosplenial interactions.

## Results

### CA1 retrosplenial-projecting (CA1-RP) neurons are a distinct class of GABAergic neurons

To characterize the presynaptic neurons in the hypothesized CA1→RSC circuit, we first anatomically localized them by injecting retrograde tracers into the RSC of wild-type (WT) mice and imaged hippocampal slices (**Fig. 1A,B**). Labeled neurons were located in dorsal CA1 at the border of stratum radiatum (SR) and stratum lacunosum-moleculare (SLM) (**Fig. 1B,C**), similar to observations in rats ^7, 8^. In agreement with prior immunostaining evidence for GABA/GAD65 expression ^7, 8^, we observed 100% co-labeling of retrograde tracer and mCherry in SR-SLM cells in Gad2-mCherry mice (double-labeling in 62 of 62 tracer-labeled cells, 6 dorsal hippocampal fields of view, 2 mice), a line with widespread expression in GABAergic neurons ^16^ (**Fig. 1D**).

**Fig. 1.**
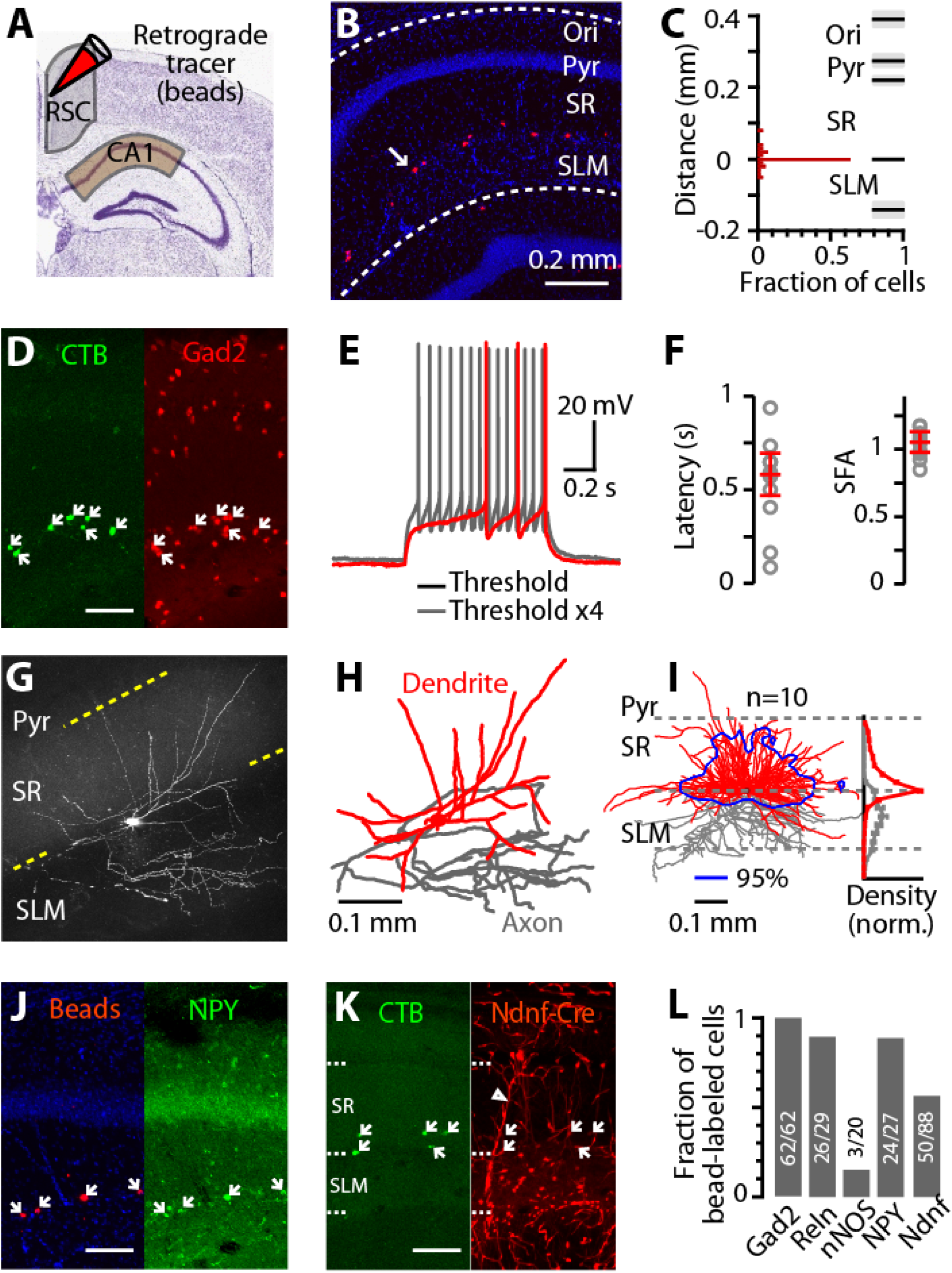
CA1 retrosplenial-projecting (CA1-RP) neurons are a distinct class of GABAergic neurons. **(A)** Schematic depicting injection of tracer into retrosplenial cortex (RSC) to label RSC-projecting neurons in dorsal CA1. **(B)** Retrogradely labeled neurons (red) in CA1 after tracer injection in RSC (one labeled neuron indicated with arrow). The slice was also labeled with DAPI (blue) to identify layers; Ori: oriens, Pyr: pyramidale, SR: radiatum, SLM: lacnosum-moleculare. **(C)** Histogram of the laminar distribution of CA1-RP neurons (2 mice, 3 and 4 slices each) relative to the SR-SLM border. Bar: 10 μm bin. **(D)** Example image of CA1 showing retrogradely labeled RSC-projecting neurons (with CTB, green) and genetically labeled Gad2-positive neurons (with mCherry, red). White arrows indicate neurons co-labeled with CTB. Scale bar: 0.1 mm. **(E)** Typical spiking pattern exhibited by CA1-RP neurons at rheobase (red) and above rheobase (gray). **(F)** Left: Plot of latency to first spike, from start of current injection step, at rheobase. Right: Plot of spike frequency adaptation ratio, measured for the first trace with ≥5 spikes as the ratio of the 4^th^/1^st^ interspike interval. Error bars: mean ± s.e.m. **(G)** Example 2-photon image (maximum intensity projection) of CA1-RP neurons (slice thickness: 300 μm). See panel H for scale bar. **(H)** Reconstructed dendritic (red) and local axonal (gray) morphology of the neuron in panel G. **(I)** Reconstructed dendrites from ten CA1-RP neurons were overlaid and aligned. Blue contour: 95% confidence interval of dendritic territory. Right: laminar profiles of dendritic (red) and axonal (gray) length density (normalized to peak). **(J)** Example image of CA1 showing retrogradely labeled RSC-projecting neurons (with beads, red), and immuno-labeled NPY-positive neurons (with FITC, green). White arrows indicate neurons co-labeled with beads. Scale bar: 0.1 mm. **(K)** Example image of CA1 showing retrogradely labeled RSC-projecting neurons (CTB, green), and genetically labeled Ndnf-positive neurons (tdTomato, red). White arrows indicate neurons co-labeled with CTB. Laminar boundaries are shown with dotted lines. Hollow arrowhead indicates endothelial-like structures, which also express Ndnf ^67^. Scale bar: 0.1 mm. **(L)** Fractions of retrogradely labeled neurons co-labeled with different markers (total number of co-labeled neurons/total number of retrogradely labeled neurons).

Because cellular electrophysiological properties can help to distinguish among GABAergic cell classes, we quantified these based on whole-cell recordings from retrogradely labeled cells in brain slices (**Table S1**). Voltage responses to current steps revealed a late-spiking pattern at rheobase, and a non-adapting regular spiking pattern above rheobase (**Fig. 1E,F**). These firing patterns are often, though not uniquely, associated with neurogliaform (NG) cells ^17, 18^.

To explore this possibility, we also characterized the morphological properties of these cells, by filling them with biocytin during recordings and imaging their dendritic and local axonal arbors (**Fig. 1G**). Analysis of images and digital reconstructions revealed dendrites extending symmetrically in the horizontal dimension along the SR-SLM border, but asymmetrically in the radial dimension, with dense branching into SR but sparse branching into SLM (**Fig. 1H,I**). Conversely, local axons branched primarily in SLM, not SR (**Fig. 1H,I**). Morphologically, these neurons thus differ markedly from NG cells described in CA1, which have dendrites localized to SLM and axons that branch profusely near the soma ^17, 19^.

Because molecular expression patterns are important for defining inhibitory cell classes, we also evaluated these. Confirming Gad2 expression (**Fig. 1D**), labeled cells were also observed at the SR-SLM border following injection of a Cre-dependent retrograde virus encoding a fluorescent protein (AAVretro-FLEX-tdTomato) ^20^ into the RSC in Gad2-Cre mice ^21^ (**Fig. S1A-C**). Consistent with prior results ^7, 8^, we observed a lack of expression of parvalbumin, calbindin, calretinin, and somatostatin (**Fig. S1F-I**). CA1-RP neurons are among several types of interneuron that selectively express the transcription factor COUP-TFII ^22^, and many COUP-TFII-positive neurons in SR-SLM are positive for α-actinin and reelin in rat ^18, 22^. We found that most CA1-RP neurons were immunopositive for reelin (double-labeling in 26 of 29 tracer-labeled cells, 3 dorsal hippocampal fields of view, 2 mice) (**Fig. 1L**, **Fig. S1J**). As neuropeptide Y (NPY) and neuronal nitric oxide synthase (nNOS) are also important interneuron markers ^17, 18^, we tested for these and found that most CA1-RP neurons expressed NPY (double-labeling in 24 of 27 tracer-labeled cells, 3 dorsal hippocampal fields of view, 2 mice), but not nNOS (double-labeling in 3 of 20 tracer-labeled cells, 3 dorsal hippocampal fields of view, 2 mice) (**Fig. 1J,L**, **Fig. S1K**). Lastly, we considered the possibility that the CA1-RP neurons express neuron-derived neurotrophic factor (Ndnf) ^23^, as was recently postulated for CA1-RP neurons as well as other retrohippocampal interneurons based on transcriptomic analysis ^24^. Following injection of retrograde tracer into the RSC of Ndnf-Cre mice (crossed with a tdTomato reporter line, Ai14), a majority of the CA1-RP neurons were tdTomato-positive (double-labeling in 50 of 88 tracer-labeled cells, 11 dorsal hippocampal fields of view from 2 mice) (**Fig. 1K,L**).

Collectively, these data show that CA1-RP neurons have some properties associated with NG cells^17, 18, 22^, but form a long-range axonal projection and also have dendritic and axonal morphology that significantly deviate from defining features of NG cells. Our characterizations establish the CA1-RP neurons as a distinct class of GABAergic neurons, anatomically positioned to mediate direct communication from dorsal hippocampus to RSC.

### CA1-RP neurons form inhibitory synapses onto RSC-L5pyr neurons at L1 apical tuft dendrites

We next investigated the circuits formed by the CA1-RP axonal projections to the RSC. Following injection of Cre-dependent AAV5-Ef1a-DIO-hChR2-EYFP into CA1 of Gad2-Cre mice (**Fig. 2A**), labeled axons were observed in L1 of the granular RSC (**Fig. 2B, Fig. S1D,E**) ^7, 8^. As a first step in analyzing CA1-RP→RSC circuits, we used an *in vivo* approach to ask whether activation of these axons in the intact brain generates a net inhibitory effect on RSC activity, as expected if connections predominantly involve GABAergic transmission to pyramidal neurons, or other effects, suggesting more complex circuits (e.g. disinhibition). In ketamine-anesthetized mice, we stimulated the ChR2-expressing CA1-RP axons within the RSC (by delivering brief, 10-ms duration photostimuli via an optical fiber placed directly over the RSC) while recording multi-unit activity on linear probes placed in L5 of the RSC (see Methods) (**Fig. 2C,D**). Photostimulation of CA1-RP axons suppressed RSC activity, to 62.8 % of baseline levels (multi-unit activity rate exceeded the -95% confidence interval of baseline, 15 to 76 ms poststimulus; baseline activity 7.2 ± 3.5 events/s; n = 8 mice) (**Fig. 2E,F**). Thus, these *in vivo* results show that CA1-RP axons can indeed exert a net inhibitory effect on RSC activity in the intact network, and point to RSC-L5pyr neurons as candidate targets of these axons.

**Fig. 2.**
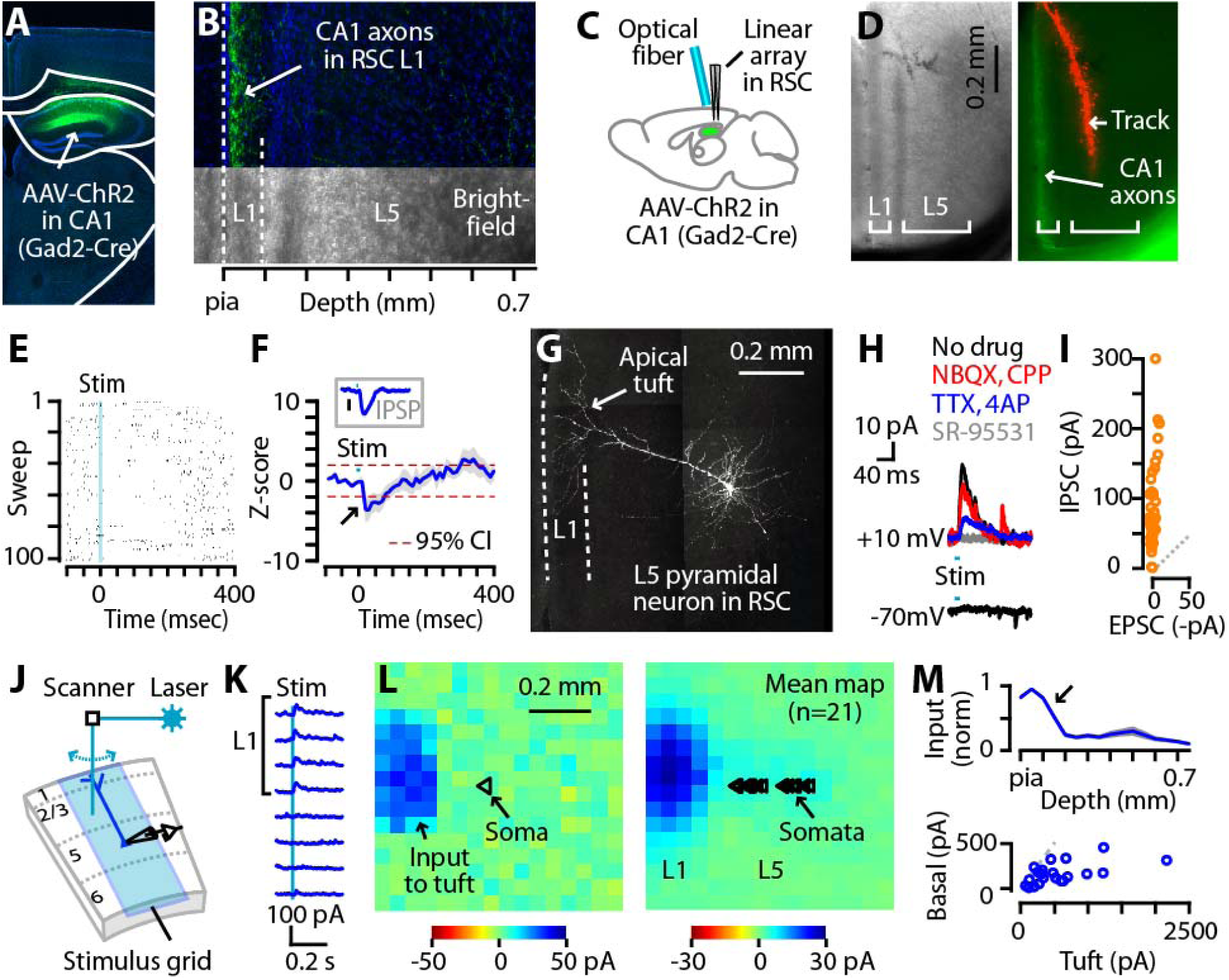
CA1-RP neurons form inhibitory synapses onto RSC-L5pyr neurons at L1 apical tuft dendrites. **(A)** Coronal section showing site of injection of Cre-dependent AAV5-Ef1a-DIO-hChR2-EYFP injection in CA1 of a Gad2-Cre mouse. Scale bar: 0.5 mm. **(B)** Bright-field (bottom) and confocal (top) images of granular RSC in a coronal section from the same CA1-injected mouse, showing L1, L5, and hChR2-expressing CA1 axons (green). **(C)** Schematic of the *in vivo* recording arrangement. **(D)** Bright-field (left) and epifluorescence (right) images of the RSC, showing CA1-RP axons in L1 and a DiI-marked electrode track in L5. **(E)** Example of peristimulus multi-unit activity recorded on one of the probe’s channels across multiple sweeps. **(F)** Plot of the averaged z-score computed from PSTHs (blue: mean; gray: ± s.e.m.; n = 8 mice; red dotted lines: 95% confidence interval; 10-ms bins). Inset: Mean IPSP waveform (blue, ± s.e.m., gray), for comparison. IPSPs, recorded in drug-free ACSF from L5pyr neurons (n = 7) in RSC slices *ex vivo*, were evoked by photostimulating CA1-RP axons in L1 of RSC with a focused laser (1 ms pulse duration). Traces were baseline-subtracted before averaging (mean resting membrane potential: -62.2 ± 1.6 mV). Bar: 0.1 mV. **(G)** 2-photon image (maximum intensity projection) of a recorded RSC-L5pyr neuron (coronal slice thickness: 250 μm), with prominent apical dendritic tuft in L1 (arrow). **(H)** Example traces showing post-synaptic currents evoked by photostimulation of CA1-RP axons at different command potentials (-70 mV for EPSC and +10 mV for IPSC) and in different pharmacological conditions. **(I)** Pairwise comparison of EPSC and IPSC recorded from multiple RSC-L5pyr neurons after photostimulation of CA1-RP axons in drug-free condition (0.1 ± 0.5 pA vs 85.2 ± 9.4 pA; p = 2.0e-08, signed-rank test). **(J)** Schematic of the sCRACM method. Whole-cell recordings were made from RSC-L5pyr neurons in slices containing ChR2-expressing CA1-RP axons, while a focused laser beam was scanned sequentially across the sites in the stimulation grid (16 x 16, 50 μm spacing) to generate an input map (see Methods). **(K)** Example peristimulus current traces from the map shown in panel H (row 7, columns 1-8). **(L)** Example (left) and mean (right) sCRACM maps of CA1-RP inhibitory synaptic to RSC-L5pyr dendrites. Triangles: soma positions of the recorded neurons. **(M)** Top: Laminar profile of CA1-RP inhibitory input (normalized to peak, mean ± s.e.m.). Bottom: Pairwise comparison of input to dendrites in L1 (left-most 3 columns) vs basal/perisomatic region (3 columns around the soma).

For detailed analysis of CA1-RP→RSC connectivity at the cellular level, we turned to *ex vivo* slice-based methods, focusing on L5Pyr neurons. These neurons, which are implicated in mediating diverse functions of the RSC^10, 25, 27^, are a prominent source of apical tuft dendrites in L1 ^28, 29^ (**Fig. 2G**). Thus, the apparent overlap in L1 of CA1-RP axons and apical tuft dendrites of L5pyr neurons implies an anatomical basis for monosynaptic GABAergic connections. However, these pre- and postsynaptic elements could in fact have minimal actual overlap, due, for example, to fine-scale micromodularities such as dendritic bundling and patchy axonal arborizations ^30^. Moreover, and more fundamentally, axo-dendritic overlap only indicates potential connectivity, and does not reliably predict actual connectivity (reviewed in ^31, 32^). We therefore used an optogenetic-electrophysiological strategy to test and characterize physiological synaptic connections from presynaptic CA1-RP axons to postsynaptic RSC-L5pyr pyramidal neurons. Recordings in acute brain slices from L5pyr neurons during wide-field photostimulation of ChR2-expressing CA1-RP axons showed outward inhibitory postsynaptic currents (IPSCs) at a command voltage of ~10 mV, but no inward excitatory postsynaptic currents (EPSCs) at -70 mV (**Fig. 2H,I**), consistent with purely GABAergic transmission. Indeed, IPSCs remained after blocking fast glutamatergic transmission (with NBQX and CPP), and after isolation of purely monosynaptic inputs (with TTX and 4-AP) ^33^, but were abolished by blocking fast GABAergic transmission (before vs after SR-95531: 31.7 ± 8.7 vs 0.3 ± 0.3 pA; n = 7, p = 0.016, signed-rank test) (**Fig. 2H**). Blocking GABA_B_ receptors caused a small reduction in IPSC amplitudes (**Fig. S2A-C**). In current clamp recordings, focal laser stimulation of CA1-RP axons in L1 (in drug-free conditions; see Methods) evoked IPSPs in RSC-L5pyr neurons that arrived and peaked with short latencies (5.2 ± 0.5 ms latency from stimulus onset to response onset, based on 10% of the peak inhibitory response; 15.8 ± 1.0 ms latency to peak inhibition; n = 7 cells), a temporal profile resembling the suppressive effect observed *in vivo* (**Fig. 2F, inset**). Thus, these results show that, at the cellular level, CA1-RP neurons form monosynaptic, GABA_A_-mediated, fast inhibitory synaptic connections to RSC-L5pyr neurons.

At the subcellular level, we wished to resolve which dendritic compartments of the L5pyr neurons receive CA1-RP input, as these can differ greatly in their computational and plasticity properties ^34, 35^. The anatomical overlap of CA1-RP axons with the apical tuft dendrites of L5pyr neurons in L1 suggests that the IPSCs might arise from inhibitory synapses at this location, despite electrotonic attenuation and imperfect space-clamp. However, a plausible alternative possibility is that the observed IPSCs could instead or additionally reflect strong perisomatic synapses. To localize the dendritic sites of CA1-RP synapses onto L5pyr neurons, we used a subcellular mapping technique, sCRACM ^33^ (**Methods, Fig. 2J**). Subcellular input maps showed that functional inhibitory synaptic contacts were localized to apical tuft dendrites in L1 (mean input to apical vs basal dendrites: 565.4 ± 108.3 vs 164.3 ± 25.4 pA; n = 21 cells, p = 8e-05, signed-rank test; **Fig. 2K-M**). These results reveal a novel apical tuft-targeting inhibitory innervation pattern, one not previously demonstrated by unbiased mapping but consistent with the anatomical axo-dendritic overlap of presynaptic CA1-RP axons in L1 and postsynaptic apical tuft dendrites of L5pyr neurons.

We performed additional studies to investigate dynamic aspects of CA1-RP→RSC-L5pyr connections. In one set of experiments, we assessed the short-term plasticity of these synapses, as activity-dependent changes in synaptic efficacy are important mechanisms involved in network dynamics ^36, 37^. To avoid potential artifacts associated with wide-field illumination of ChR2-expressing presynaptic terminals, we used an approach involving repetitive laser stimulation of axons away from postsynaptic dendrites ^38^. Trains of IPSCs evoked by focal stimulation of CA1 axons in L1 (in drug-free conditions; see **Methods** for details) displayed strong synaptic depression (**Fig. S3A,C,D**). In other experiments we found that these apical dendritic IPSPs were sufficiently potent, despite their electrotonic remoteness, to prolong inter-spike intervals during trains of action potentials in the L5pyr neuron evoked by current injection (see **Methods; Fig. S4**). These results demonstrate that, on a millisecond time scale, and consistent with the *in vivo* results presented above, activation of monosynaptic CA1-RP inhibitory input is capable of briefly suppressing activity of RSC-L5pyr neurons. However, due to synaptic depression, the strength of this inhibitory influence likely diminishes during repetitive activity.

Collectively, these analyses of CA1-RP→RSC-L5pyr circuits show that CA1-RP axons form monosynaptic GABA_A_-mediated inhibitory synapses onto RSC-L5pyr neurons, via depressing synapses made primarily onto apical dendritic tufts, capable of exerting a rapid, inhibitory effect on RSC activity.

### ATN-TC axons form excitatory synapses onto RSC-L5pyr neurons at L1 apical tuft dendrites

In addition to these direct projections from CA1, hippocampus communicates with RSC via pathways involving the anterior thalamic nuclei (ATN), particularly the anteroventral (AV) nucleus ^39, 40^, which sends a thalamocortical (TC) projection ramifying mainly in L1 of RSC ^14^ - anatomically strikingly similar to that of CA1-RP axons, but presumably having an opposite, excitatory effect. To test this, we characterized ATN-TC connectivity to RSC-L5pyr neurons, using a similar multi-step approach as for the CA1 projection.

First, we injected the granular RSC in WT mice with tracer to retrogradely label TC neurons in thalamus, which verified that AV is the major source of its thalamic afferents (**Fig. 3A**) ^15^. Then, we injected AV with AAV1-CamKIIa-hChR2-mCherry to anterogradely label TC axons, which indicated that in mice these axons indeed target primarily L1 in granular RSC, and particularly L1a (its outermost sublayer), with additional sparser branching in deeper layers and in dysgranular RSC (**Fig. 3B**). To further assess if these L1 axons in granular RSC arise from AV neurons, which may branch in other layers ^14^, and/or from TC neurons in surrounding nuclei such as the anterodorsal (AD) nucleus ^15, 41^, we repeated this experiment using Grp_KH288-Cre mice ^42^, a driver line showing Cre-expressing neurons in a subset of AV nucleus but none in AD (**Fig. S5A-H**). Following injection of AV with Cre-dependent AAV5-Ef1a-DIO-hChR2-EYFP, we again observed labeled axons in L1 (**Fig. 3C,D**), supporting the results obtained with WT mice indicating that AV, rather than AD, is the main source of L1-targeting projections from the ATN to the granular RSC. Since targeted injections into AV in WT mice resulted in more extensive labeling of AV-TC neurons (than in the Cre line), we used this approach for further study of these AV-predominant ATN→RSC projections.

**Fig. 3.**
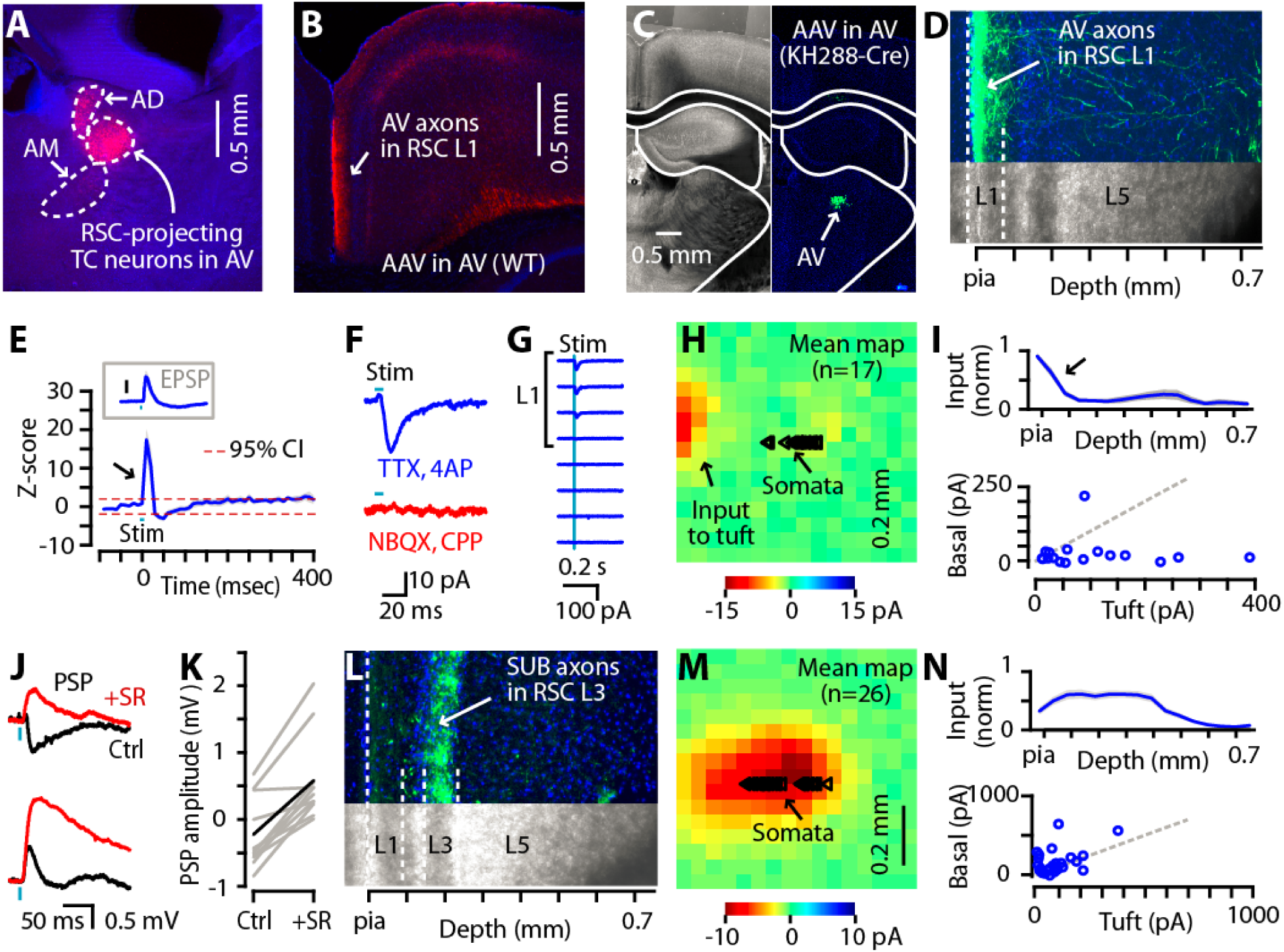
ATN-TC axons make excitatory synapses onto RSC-L5pyr neurons at L1 apical tuft dendrites. **(A)** Coronal section showing labeling in AV and AD after retrograde CTB647 injection into RSC. **(B)** Axonal projection pattern in RSC resulting from AAV1-CamKIIa-hChR2-mCherry injection targeted to AV (coronal section). **(C)** Labeling in AV (Grp_KH288-Cre mouse) at the site of injection of AAV5-Ef1a-DIO-hChR2-EYFP. **(D)** Axonal projection to RSC resulting from the injection shown in panel C. **(E)** Plot showing averaged z-score computed from PSTH (blue). Dotted lines (red) indicate 95% confidence interval. Bin: 10 ms. Inset: Mean EPSP waveform (blue, ± s.e.m., gray), for comparison, recorded in drug-free ACSF from L5pyr neurons (n = 7) in RSC slices *ex vivo*, while focally photostimulating ATN-TC axons in L1 of RSC with a focused laser. Traces were baseline-subtracted before averaging (mean resting membrane potential: -64.3 ± 2.8 mV). Bar: 0.5 mV. **(F)** Example EPSC recorded in a RSC-L5pyr neuron after photostimulation of ATN-TC axons in different pharmacological conditions. **(G)** Example peristimulus current traces extracted from one neuron’s sCRACM map. **(H)** Mean sCRACM map of ATN-TC excitatory synaptic input to RSC-L5pyr dendrites. Triangles: soma positions of the recorded neurons. **(I)** Top: Laminar profile of ATN-TC excitatory input (normalized to peak, mean ± s.e.m.). Bottom: Pairwise comparison of input to dendrites in L1 (left-most 3 columns) vs basal/perisomatic region (3 columns around the soma). (**J**) Example traces of photo-evoked PSPs recorded from two different L5pyr neurons, before (black) and after (red) application of SR-95531 to block CA1-RP inputs. In one neuron (top), SR-95531 changed the photo-evoked response from net inhibitory to excitatory; in another neuron (bottom), it enhanced the size of EPSP. (**K**) Comparison of PSP amplitude (mean of 30 ms from stim onset) before and after SR-95531. Grey lines are for individual recordings, and black indicates the group mean. (**L**) Axonal projection pattern in RSC resulting from injection of AAV1-CamKIIa-hChR2-mCherry into dorsal subiculum. (**M**) Mean sCRACM map of subicular excitatory synaptic input to RSC-L5pyr dendrites. Triangles: soma positions of the recorded neurons. (**N**) Same as panel H, but for subicular inputs.

Next, we assessed the impact of ATN-TC input on RSC activity in the intact brain using the same *in vivo* approach as for the CA1-RP inputs (see above, and **Methods**), recording with linear arrays to sample multi-unit activity in L5 of RSC while stimulating ChR2-expressing TC axons (labeled by injecting AAV-ChR2 in WT mice) by delivering brief photostimuli via an optical fiber placed directly over the RSC. Photostimulation of ATN-TC axons rapidly and sharply increased RSC activity above baseline levels (multi-unit activity rate exceeded the +95% C.I. of baseline, ~3 to 27 ms poststimulus; baseline activity 7.5 ± 2.8 events/s; n = 6 mice) (**Fig. 3E**), followed by suppression (activity exceeded the -95% C.I. of baseline, ~31 to 58 ms poststimulus). Thus, these *in vivo* results show that ATN-TC axons can indeed exert a brief, net excitatory effect on RSC activity in the intact network, and point to RSC-L5pyr neurons as candidate targets of these axons.

We then assessed ATN→RSC synaptic connectivity at the cellular level, using slice-based methods as for the CA1-RP connections (see above, and **Methods**). Wide-field photostimulation of ATN-TC axons in RSC slices generated EPSCs in L5pyr neurons, sampled at a command voltage of -70 mV. At command voltage of ~10 mV, short-latency IPSCs could also be detected, suggesting feedforward (e.g. disynaptic) inhibition (**Fig. S5I,J**). Confirming this, only the inward postsynaptic current remained after bath application of TTX and 4-AP (**Fig. S5I**), indicating that the EPSCs are monosynaptic and the IPSCs are disynaptic. Application of blockers of glutamatergic transmission (NBQX and CPP) abolished the EPSCs (before vs after drugs: -7.1 ± 2.0 vs -0.5 ± 0.4 pA; n = 5 cells, p = 0.01, 2-tailed paired t-test), indicating that ATN-TC axons provide monosynaptic glutamatergic transmission to L5pyr neurons (**Fig. 3F**). In current clamp recordings, focal laser stimulation of ATN-TC axons in L1 evoked EPSPs in RSC-L5pyr neurons that arrived and peaked with short latencies (4.5 ± 0.4 ms latency from stimulus onset to response onset, based on 10% of the peak excitatory response; 9.0 ± 1.3 ms latency to peak excitation; n = 7 cells), a temporal profile resembling the excitatory effect observed *in vivo* (**Fig. 3E, inset**). Thus, these results show that, at the cellular level, ATN-TC neurons form monosynaptic, glutamatergic, excitatory synaptic connections to RSC-L5pyr neurons.

We further analyzed ATN-TC→RSC-L5pyr connections to localize the dendritic sites of input and assess dynamic properties. Mapping the subcellular location of ATN-TC inputs (by sCRACM; see above, and **Methods**) showed that, on average, ATN-TC inputs targeted the apical tuft dendrites of L5pyr neurons (mean input to apical vs basal dendrites: -103.2 ± 25.5 vs -26.2 ± 12.5 pA; n = 17 cells, p = 0.002, signed-rank test; **Fig. 3G-I**). In separate experiments, evaluating the short-term plasticity of these synapses (using repetitive laser stimulation aimed at the ATN axons in L1; see above, and **Methods**) showed that trains of EPSCs displayed a pattern of non-depressing initial responses followed by depressing later responses (**Fig. S3B,C,D**). In other experiments, current-clamp recordings from RSC-projecting AV neurons showed that these displayed expected patterns of membrane potential-dependent burst-firing ^43, 44^ (**Fig. S5K**). Results from these experiments thus add important information about synaptic localization and dynamic signaling in ATN-TC→RSC-L5pyr circuits.

Given that CA1-RP and ATN-TC axons both target L1, and both monosynaptically innervate L5pyr neurons at their apical tuft dendrites, do individual RSC-L5pyr neurons receive both ATN-TC and RSC-RP inputs? To test this, we co-stimulated ChR2-expressing CA1 and ATN axons in L1 and measured post-synaptic potentials (PSPs) before and after blocking CA1 inputs with SR-95531, with TTX/4-AP in the bath and Cs^+^-based internal solution (see **Methods** for details). Overall, the photo-evoked PSP became significantly more positive after removal of CA1 inputs (before vs after 10 μM SR-95531: -0.20 ± 0.17 vs 0.60 ± 0.21 mV; n = 10 cells, p = 0.002, signed-rank test) (**Fig. 3J,K**). These data indicate that the two types of input tend to target a common pool of RSC L5 neurons in a convergent manner, rather than forming parallel pathways innervating separate pools.

Another source of excitation to RSC-L5pyr neurons is from pyramidal neurons in subiculum ^13^. We used sCRACM to map the dendritic locations of these inputs, since this can help to understand how postsynaptic mechanisms integrate all three inputs (CA1, ATN, and subiculum). Subicular inputs mainly innervated basal dendrites of L5pyr neurons, with relatively weak input to apical tuft dendrites in L1 (mean input to apical vs basal dendrites: -84.9 ± 16.9 vs -157.2 ± 31.2 pA; n = 26 cells, p = 0.016, signed-rank test; **Fig. 3M,N**). In separate experiments we characterized the short-term plasticity of these synapses, which showed a pattern of facilitating responses (**Fig. S5B,C,D**). We previously characterized the spiking patterns of the presynaptic RSC-projecting neurons in the subiculum as burst-firing ^13^. These results indicate that subicular axons target a more proximal subcellular dendritic compartment and express a facilitating pattern of short-term plasticity compared to CA1 and ATN inputs to L5pyr neurons.

Overall, these findings show that ATN-TC axons form monosynaptic glutamatergic connections onto RSC-L5pyr neurons that target apical tuft dendrites and can exert a rapid, excitatory effect on RSC activity. Together with the previous results, this indicates that ATN-TC projections and CA1-RP projections form a convergent triadic circuit with RSC-L5pyr neurons, with anatomical alignment of afferent input at the cellular and subcellular levels, and opposing actions.

### Opposing actions of CA1-RP inhibition and ATN-TC excitation in contextual fear learning

The anatomical convergence of CA1-RP→RSC and ATN-TC→RSC projections onto the apical tuft dendrites of L5pyr neurons combined with the physiologically opposing actions of neurotransmission suggests the hypothesis that these circuit connections contribute differentially to the functions of the RSC in processing hippocampal-dependent memories. To explore this, we used contextual fear conditioning (CFC), a behavioral paradigm that induces hippocampal-dependent contextual fear memory in mice and modulates RSC activity ^45-50s^.

To test the roles of each pathway in memory encoding, we selectively silenced presynaptic axons from either source at their arborizations within the RSC during CFC, using a previously established chemogenetic approach based on expressing hM4D(Gi) in presynaptic neurons and blocking synaptic release from a subset of axonal branches by local infusion of clozapine-N-oxide (CNO) ^51^ (**Methods**; **Fig. 4A**). As in prior studies using this approach ^13^ we performed control experiments to test the efficacy of the technique, which confirmed that CNO application reduced synaptic transmission in both circuits (**Fig. S6**).

**Fig. 4.**
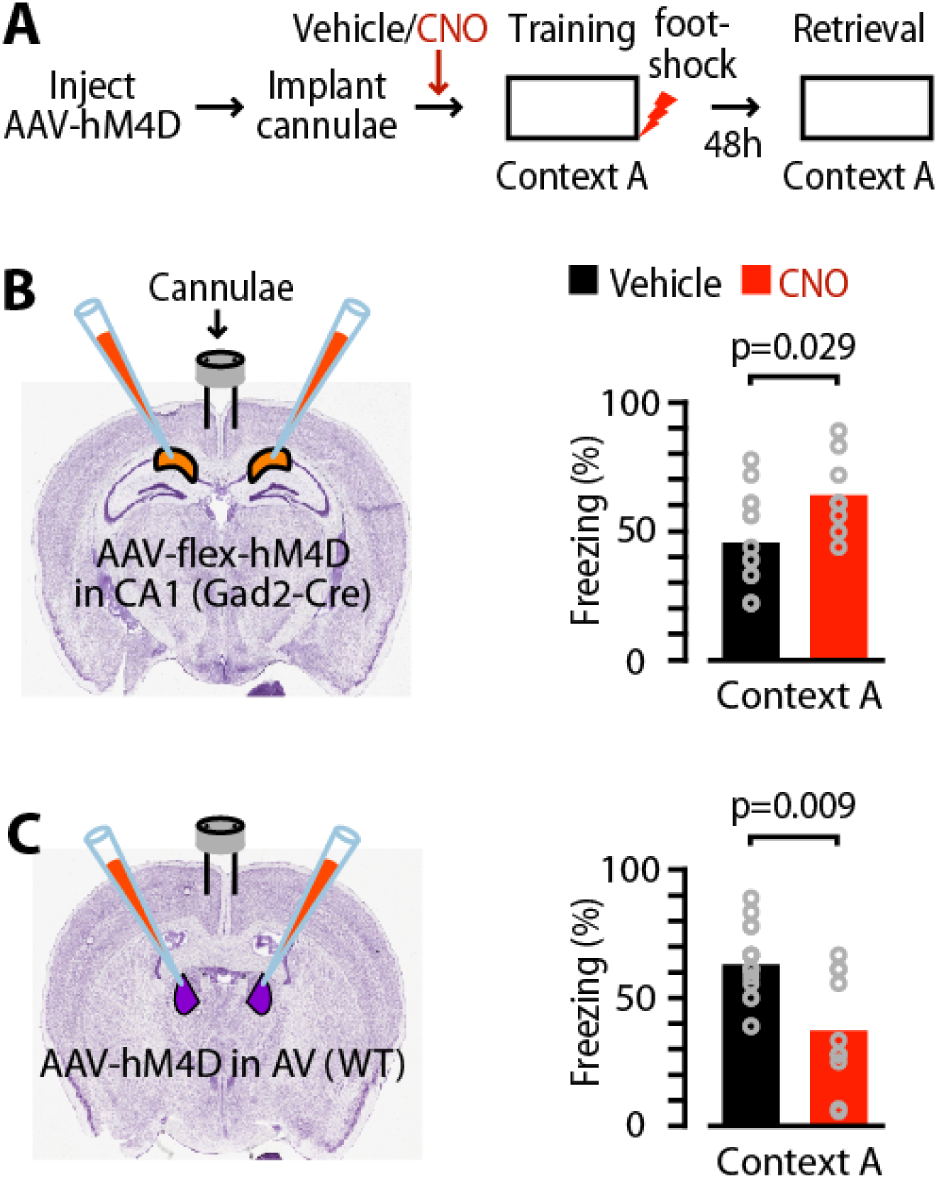
Opposing actions of CA1-RP inhibition and ATN-TC excitation in contextual fear learning. **(A)** Schematic of the experimental paradigm. **(B)** Left: Schematic illustrating bilateral injection of AAV8-hSyn-DIO-hM4D(Gi)-mCherry into CA1 of Gad2-cre mice and placement of cannulae in RSC for CNO/ACSF infusion during CFC. Right: Freezing score (%) during exposure of mice (n = 14 and 9 mice for vehicle and CNO group, respectively) to conditioned context during retrieval testing. **(C)** Left: Schematic illustrating bilateral injection of AAV8-hSyn-HA-hM4D(Gi)-mCherry into AV of WT mice. Right: same as B, but for assessment of ATN→RSC connections in CFC (n = 17 and 10 mice for vehicle and CNO group, respectively).

Mice underwent bilateral injections of virus (Cre-dependent or -independent AAV-hM4D(Gi), as appropriate) into either CA1 (Gad2-Cre mice) or AV (WT mice), followed by bilateral implantation of cannulae in RSC, which were used to infuse either CNO (CNO group) or vehicle (vehicle group) 30 min before CFC. We have previously shown that CNO infusion into the RSC of wild-type mice (i.e., without hM4D(Gi) expression) did not induce off-target effects during CFC ^13^. In additional control experiments, adding a fluorescent dye to the infusant showed that infusions remained in the RSC, without spread to subiculum or CA1 (**Fig. S6J**). Silencing of CA1-RP axons in the RSC during CFC increased freezing behavior during the retrieval test in the CNO-treated group compared to controls (mean freezing, vehicle vs CNO group: 45.6 ± 5.0% vs 64.1 ± 4.9%; n = 14 vs 9 mice, p = 0.03, rank-sum test; **Fig. 4B**). CNO did not affect activity in the conditioning box during training or activity burst to the shock (mean activity during training, vehicle vs CNO group: 9.5 ± 1.9 cm/s vs 11.8 ± 2.5 cm/s; p = 0.46, rank-sum test; mean activity burst to the shock: 25.6 ± 1.9 cm/s vs 30.3 ± 8.4 cm/s; p = 0.69, rank-sum test). In contrast, silencing of ATN-TC axons in the RSC during CFC caused a significant reduction in freezing behavior in retrieval test in the CNO-treated group compared to controls (mean freezing, vehicle vs CNO group: 62.9 ± 3.4% vs 37.1 ± 7.2%; n = 17 vs 10 mice, p = 0.009, rank-sum test; **Fig. 4C**). Again, CNO did not affect activity in the conditioning box during training or activity burst to the shock (mean activity during training, vehicle vs CNO group: 15.4 ± 1.2 cm/s vs 19.5 ± 2.0 cm/s; p = 0.08, rank-sum test; mean activity burst to the shock: 57.7 ± 2.8 cm/s vs 59.0 ± 4.5 cm/s; p = 0.86, rank-sum test). These data indicate that CA1-RP→RSC and ATN-TC→RSC circuits are both engaged during the encoding of contextual fear memory, but appear to serve opposing roles, with the inhibitory pathway normally suppressing and the excitatory pathway enhancing the expression of context memories.

## Discussion

Harnessing the availability of an array of tools for targeted analysis of long-range circuits in the mouse, we dissected the synaptic connectivity mediating the confluence and interaction of major afferent projections to the RSC from dorsal hippocampus and ATN. Our findings provide cellular- and subcellular-level mechanistic insight into information-processing in these hippocampo-thalamo-cortical networks (**Fig. 5**).

**Fig. 5.**
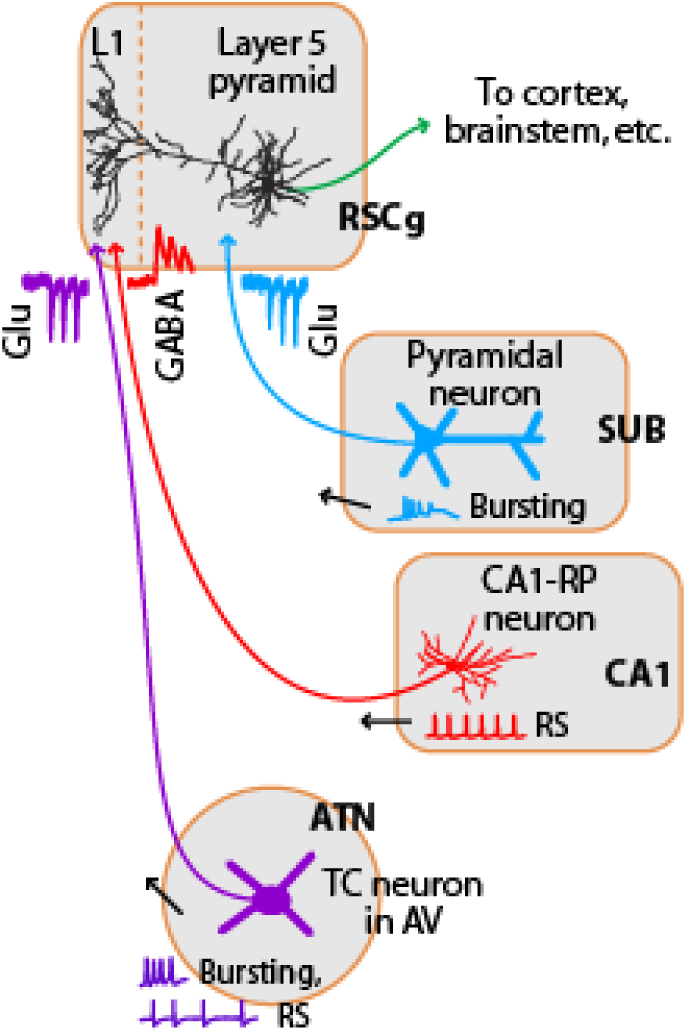
Cellular and subcellular organization of long-range input circuits of the RSC. Inhibitory axons from CA1-RP neurons and excitatory axons from ATN thalamocortical neurons, particularly in the AV nucleus, converge in L1 of RSC and innervate the distal, apical-tuft dendrites of L5pyr neurons in the granular RSC, with opposing neurotransmitter actions. Excitatory axons from dorsal subiculum innervate more proximal dendrites in deeper layers. Activity-dependent synaptic properties (traces near the synaptic connections) and somatic firing patterns (traces near presynaptic somata) are distinct for each source of afferent input. RS: regular-spiking.

Our characterizations of CA1-RP neurons, the source of the CA1 inhibitory projection to RSC, reveal a unique combination of properties defining them as a distinct GABAergic cell class. Certain properties, such as firing patterns and molecular expression, resemble those described in various types of local interneurons, particularly NG cells. However, they exhibit unusual features not shared with NG cells, including asymmetric dendritic morphology and mostly GABA_B_-independent synaptic transmission. In addition, although their axons innervated apical tuft dendrites in L1 like local NG cells ^19, 52^, they did so through long-range projections. We also observed variability in marker labeling (e.g. Ndnf), suggesting some degree of within-class heterogeneity. The results of our characterizations appear consistent with a recent single-cell transcriptomic analysis that assigns retrohippocampally projecting CA1 interneurons, including putative CA1-RP neurons, to a cluster of cells – the Ntng1.Rgs10 subgroup in “continent 6” – that form long-range inhibitory projections and lack expression of most classical interneuron markers ^24^. Our results thus help to define the cellular identity and basic characteristics of these long-range-projecting, apical tuft-targeting, CA1-RP inhibitory neurons.

Optogenetic-electrophysiological circuit analyses showed that CA1-RP axons directly inhibited the apical tuft dendrites of RSC-L5pyr neurons, converging there with excitatory connections from ATN-TC neurons, while excitatory subicular inputs mainly innervate the perisomatic dendrites in deeper layers (**Fig. 5**). This circuit configuration bears close resemblance to generalized cortical circuit models in which L5pyr neurons receive dual excitatory inputs to their apical tuft dendrites via L1 (e.g. from matrix thalamus) and to proximal dendrites via deeper layers (e.g. corticocortical), which synergistically drive firing ^34, 53^. The essential concept is that when both input channels are active they are associatively paired, through active dendritic conductances, resulting in robust firing of the postsynaptic pyramidal neuron ^34^. Similar circuits and pairing mechanisms have been identified in hippocampus, where coincident activation of perforant-path and CA3 inputs to apical and proximal dendrites of CA1 pyramidal neurons drives non-linear responses and synaptic plasticity ^54^. Importantly, the associative pairing mechanism can be suppressed by local apical tuft-targeting interneurons ^34^. The RSC circuits delineated here appear remarkably analogous – with the extraordinary variation that the apical tuft inhibition is supplied remotely, by CA1-RP neurons. To the extent that the universal model proposed by Larkum ^34^ holds for these RSC circuits, a clear implication is that one function of CA1-RP→RSC inhibition is to suppress the associative pairing, by RSC-L5pyr neurons, of information carried by thalamic and subicular afferents.

CA1-RP→RSC circuits are likely to have multiple other functional roles as well. Some are suggested by analogy to functions ascribed to local apical tuft-targeting interneurons in neocortex ^55^, including modulation of sensory input-driven dendritic events ^52, 56^, and the promotion of “robustness” by reducing the variability of cortical neuronal responses to sensory inputs ^57^. In this way, CA1-RP inhibition might contribute to the “sharpening” of contextual memory representations ^58^. By analogy to GABAergic projections from zona incerta to neocortex, which similarly ramify in L1 and inhibit L5pyr neurons ^3^, CA1-RP neurons might be important for cortical development, or for braking excitation to prevent epileptiform activity. They may be involved in oscillations and rhythmogenesis, by analogy to long-range inhibitory projections in septo-hippocampal networks ^59^. The behavioral roles of CA1-RP neurons’ circuits may be similarly protean. Chemogenetic disconnection of the inhibitory CA1 or excitatory ATN inputs to RSC gave opposing effects on one aspect of behavior, the encoding of fear memory. However, while this accords with the opposing neurotransmitter actions of these pathways, the contributions of these circuits to other aspects of RSC-dependent behavior are likely more complex. Nevertheless, our results, together with our recent findings demonstrating contributions of excitatory subiculum→RSC pathways to contextual fear conditioning ^13^, suggest a role for the inhibitory CA1-RP→RSC projection in modulating the impact of thalamic inputs relative to that of subicular inputs, perhaps to avoid inappropriate and potentially pathological behavioral responses due to unregulated associative pairing of thalamic and subicular inputs. Indeed, dysfunction in RSC-related pathways has been implicated in a variety of neurological and psychiatric disorders ^11^, as has dysfunction in TC circuits ^39, 60^. Our findings thus additionally present a detailed framework (**Fig. 5**) for investigating CA1-RP neurons and RSC circuits as etiological factors in these conditions, and as targetable cellular candidates for novel therapeutic interventions.

## Materials and Methods

**Animal care and mouse lines.** Studies were approved by Northwestern University Animal Care and Use Committee and followed the animal welfare guidelines of the National Institutes of Health. All experiments were performed using mouse strain C57BL/6 or transgenic mice with C57BL/6 background (male or female). Mice were 9-20 weeks old at the time of electrophysiological and behavioral experiments. Mice were kept on a 12-hour light/dark cycle and had unrestricted access to the food and water.

Gad2-Cre (RRID:IMSR_JAX:010802) ^21^. Gad2-mCherry (RRID:IMSR_JAX:023140) ^16^, and Ndnf-Cre (RRID:IMSR_JAX:028536) ^23^ mice were obtained from Jackson Laboratory. Grp_KH288-Cre mice (RRID:MMRRC_037585-UCD) ^42^ were obtained from MMRRC, and back-crossed in-house with C57BL/6 mice for at least 6 generations. All Cre lines were maintained by crossing with C57BL/6 mice. Experiments were performed using heterozygous mice, identified to be positive for Cre with genotyping. Gad2-mCherry mice were maintained by homozygous breeding. In some cases, heterozygous litters of each Cre lines were crossed with tdTomato reporter line Ai14 from Jackson laboratory (RRID:IMSR_JAX:007908).

**Viral vectors.** For Cre-dependent anterograde expression of ChR2 and hM4D(Gi), AAV5-Ef1a-DIO-hChR2(E123T/T159C)-EYFP-WPRE-hGH (Addgene 35509) and AAV8-hSyn-DIO-hM4D(Gi)-mCherry (Addgene 44362) were used. For Cre-independent anterograde expression of ChR2 and hM4D(Gi), AAV1-CamKIIa-hChR2(E123T/T159C)-mCherry-WPRE-hGH (Addgene 35512) and AAV8-hSyn-HA-hM4D(Gi)-mCherry (Addgene 50475) were used. For Cre-dependent retrograde expression, AAVrg-CAG-Flex-tdTomato-WPRE (Addgene 51503) was used. Viral vectors were purchased from University of Pennsylvania Viral Vector Core or Addgene.

***In vivo* stereotaxic injections.** Mice were anesthetized with isoflurane, head-fixed to a stereotaxic frame, and thermally supported with a feedback-controlled heating pad (DC Temperature Control System, FHC). Buprenorphine (0.3 mg/kg) and meloxicam (1 mg/kg) were injected subcutaneously for post-operative pain relief. After incising the scalp over the cranium, small craniotomy was opened using a dental drill over the RSC, dorsal CA1 of the hippocampus, or the AV nucleus of the thalamus. The stereotaxic coordinates of the RSC target were (relative to bregma, in mm) anteroposterior -1.6 and 2.0, lateral 0.1, and ventral 0.3, 0.6 and 0.9, CA1 target were anteroposterior -2.0, lateral 1.5, and ventral 1.4, and those of the AV target were anteroposterior -0.5, lateral 1.2, and ventral 2.8. A beveled injection pipette, back-filled with mineral oil and front-filled with AAV solution, was slowly advanced to the target depth, where a small volume (50 nL) of virus was injected using a displacement-driven injector (MO-10 Narishige). The pipette was left in place for 5 mins before retraction. Once retracted, the incision was closed with a nylon or silk suture.

**Slice electrophysiology.** Mice were euthanized 3-5 weeks after virus injection and coronal brain slices (250 µm thick for RSC and AV, 300 µm thick for CA1) were prepared using a vibratome (VT1200S, Leica) in ice-cold choline-based cutting solution (composition, in mM: 25 NaHCO_3_, 1.25 NaH_2_PO_3,_ 2.5 KCl, 0.5 CaCl_2_, 7 MgCl_2_, 110 choline chloride, 11.6 sodium L-ascorbate, and 3.1 sodium pyruvate). The rostral end of the brain was first removed by a coronal blocking cut. The remaining part of the brain was mounted caudal-end-up on the exposed surface. Slices were collected in a caudal-to-rostral manner, starting with the first slice in which the corpus callosum was observed and proceeding for four more slices, spanning the granular RSC. Recordings were mostly made from the middle three slices in this series. Slices were transferred to artificial cerebrospinal fluid (ACSF; composition, in mM: 127 NaCl, 25 D-glucose, 2.5 KCl, 1 MgCl_2_, 2 CaCl_2_, and 1.25 NaH_2_PO_3_) for 30 minutes at 34 ^o^C and then 1 hour or more at room temperature (~21 ^o^C) before recording. Whole-cell recordings were performed using an upright microscope (BX51WI, Olympus) equipped with gradient-contrast and epifluorescence optics. Pipettes (~2.5-4 MΩ) were filled with either cesium- or potassium-based internal solution containing (in mM): 128 potassium or cesium methanesulfonate, 10 HEPES, 10 phosphocreatine, 4 MgCl_2_, 4 ATP, 0.4 GTP, and 3 ascorbate, pH 7.25, 290-295 mOsm. For cesium-based solution, 1 mM QX-314 and 1 mM EGTA was also included. To visualize neurons after the recording, 0.05 mM Alexa 647 hydrazide and/or biocytin (4 mg/mL) was added to internal solution. Using a 60× objective lens (LUMPlanFI/IR, N/A 0.9, Olympus), recordings were targeted either to L5pyr neurons (RSC slices) or fluorescently labeled RSC-projecting neurons in CA1 (hippocampal slices). ACSF, oxygenated with 95% O_2_/5% CO_2_, was perfused using a pump-driven recirculation system, and the recording temperature was maintained at 32 ^o^C (most experiments) or 34 ^o^C (intrinsic-property recordings) using an in-line temperature controller (TC-324B, Warner instrument). Data were acquired using Ephus software ^61^. Signals were amplified with an Axon Multiclamp 700B (Molecular Devices), filtered at 4 kHz, and sampled at 10 kHz (most experiments) or 40 kHz (intrinsic-property recordings).

**Wide-field ChR2 photostimulation.** Wide-field photostimulation was performed as described previously ^62^. Brief (5 ms) flashes from a blue LED (1 mm^2^/mW, M470L3, Thorlab) were delivered through a 4× objective lens (UPlanSApo, N/A 0.16, Olympus) focused onto the specimen, by gating the LED output with a TTL signal. Trials were repeated at 10 s intervals. Recordings were made in voltage-clamp mode and drug-free conditions, with the command potential set to -70 mV to record EPSCs. It was then changed to 0-10 mV to record IPSCs from same neuron. Tetrodotoxin (TTX, 1 µM) and 4-aminopyridine (4-AP, 100 µM) were added to isolate monosynaptic connections ^33^. Traces were analyzed offline by averaging 3-5 trials per condition, baseline-subtracting based on the 100 ms pre-stimulus, and calculating the mean amplitude over the time window 0 to 50 ms from stimulus onset.

**Subcellular ChR2-assisted circuit mapping (sCRACM).** sCRACM was performed as described previously ^33, 63^ using a laser scanning photostimulation system consisting of a blue laser (model MLL-FN-473, 473 nm, 50 mW, CNI Laser), electro-optical modulator (model 350-50, Conoptics), mechanical shutter (model VMM-D1, Uniblitz), and mirror galvanometers (model 6210, Cambridge Technologies). The laser beam was focused through a 4× objective lens onto the slice (~60 µm diameter, full-width at half maximum). Laser power was monitored using a photodiode (part number 53-379, silicon detector, blue enhanced response, 100 mm^2^, Edmund Optics) and adjusted by a graded neutral density filter to 0.75 mW at the specimen. An image of the slice was captured using the 4× objective lens, oriented with the pia up, and overlaid with a graphical representation of the photostimulation grid (16 by 16 square array with 50 µm spacing), which was rotated and offset to align its top row with the pia and its middle columns with the soma. Each site was stimulated in pseudo-random order at an inter-stimulus interval of 0.4 or 1 s. Mapping was repeated 3 times for each neuron, using a different pseudo-random sequence each time. In experiments involving GABA_B_ pharmacology (**Fig. S2**) and current-clamp recordings (insets of **Fig. 2M, 3I**), the laser scanning system was used to photostimulate axons at a single spot in L1.

**Repetitive stimulation of axons.** Brain slices were prepared from CA1-, AV-, or subiculum-injected mice, as described above. Whole-cell recordings were performed in voltage-clamp mode with Cs^+^-based internal solution, in drug-free conditions (plain ACSF, without TTX/4-AP), as described above. Sites in L1 (for CA1 and ATN axons) or deeper layers (L3/5, for subicular axons), approximately ~250 um lateral to either side of soma, were selected for targeted focal photostimulation, to avoid potential artifacts associated with direct stimulation of ChR2-expressing axon terminals (“over-bouton stimulation”) ^38^. Stimulus intensity was set to ~0.75 mW at the focal plane. The laser beam was flashed (1 ms duration) at the selected sites in a sequential manner (1 s inter-stimulus interval), to identify a site of input. If no responses were observed, new sites were tested. Once a suitable hotspot of input was located, axons at this site were repetitively stimulated with a train of 10 pulses (1-ms duration each), at 20 Hz. Ten sweeps were collected, and traces were analyzed offline to characterize mean response amplitudes and short-term dynamics.

***Ex vivo* analysis of convergent inputs.** In this case, both CA1 and AV were injected (with Cre-dependent and Cre-independent versions of AAV-ChR2, respectively, as described above) in the same animals (Gad2-Cre mice), to express ChR2 in both input pathways to the RSC. Methods were as described above for each input pathway separately. Optogenetic photostimulation and electrophysiological recordings were made in slices as described above, with TTX/4-AP in the bath solution to prevent disynaptic responses. The laser beam was targeted to L1, and brief (1 ms) photostimuli were delivered to activate ChR2-expressing axons and their presynaptic terminals synapsing onto the recorded L5pyr neurons. After sampling pre-drug responses under control conditions, SR-95531 was added to the bath (10 μM), and post-drug responses were collected.

**2-photon microscopy.** Slices with biocytin-filled neurons were fixed in 4% PFA (in 0.1M PBS) overnight, then washed in PBS before incubating in PBS containing 2% Triton X-100 (Sigma) and streptavidin conjugated with Alexa568 (1:200, Invitrogen) for at least 14 hours at 4 ^o^C. Slices were washed in PBS the next day and mounted onto a glass coverslip (#1.5, 22 x 40 mm, Warner Instruments), imaging side down. A well for the slice was crafted from two 1.5 mm thick glass coverslip (#1, 22 x 22 mm, Warner), which were stacked and attached to the short edges of the main coverslip. After removing excess PBS from the slice, drops of mounting media (FluorSave, Millipore) were applied before sealing with glass slide (75 x 25 x 1 mm, Ever Scientific). Once the mounting media solidified, the slide was sealed with nail polish.

Two-photon imaging was performed on a Nikon A1R MP+ Multiphoton system using a water-immersion 25× objective lens (APO LWD, N/A 1.10, WD 2.0 mm, optimized for coverglass thickness 0.17 mm, Nikon) with the laser (Ti:sapphire, Chameleon Vision, Coherent) tuned to 860 nm wavelength. Image stacks were acquired using Nikon NIS-Elements software, with settings of at 1024 x 1024 pixels (0.5 μm x-y pixel size), 2.4 μs dwell time, and 2 μm z-step size. The laser power and the gain of the GaAsP photomultiplier detectors were depth-adjusted.

**Morphological reconstructions.** Morphologies of CA1 neurons were digitally 3D-reconstructed in a semi-automated manner with Imaris software (Bitplane, Inc.), using the AutoPath feature in the Filament Tracer module. Reconstructions were exported in Imaris format, converted into ASCII or SWT format using NLMorphologyViewer_software (version 0.4.0.dev_x86.msi; NeuronLand, http://neuronland.org) and Matlab functions, and further analyzed using Matlab routines to align them and quantify length density ^64^. Length-density maps of individual neurons were pooled and averaged. A contour plot of the resulting average map was used to determine the average 95% contour level.

**Immunohistochemistry.** For immunostaining of CA1 slices, mice were first injected in the RSC with retrograde tracer (~50 -100 nL), either red RetroBeads (undiluted; Lumafluor) or cholera toxin subunit B conjugated with Alexa647 (CTB647, 100 mg in 20 μL PBS; Invitrogen). After two or more days, to allow for retrograde transport, mice were transcardially perfused with ice-cold PBS (0.1M, pH 7.4) followed by 4% PFA (w/v in PBS). Brains were extracted and incubated in the same PFA solution overnight at 4 ^o^C. The brains were washed with PBS, and sliced (in PBS) into 100 μm sections using a microtome (HV650 V, Microme). In some cases (SOM and CR), brains were cryoprotected in PBS containing sucrose (20% then 30%, w/v), and then frozen in OCT compound (Fisher), before being cut into 50 μm section using a cryostat. Selected sections were transferred to a 24-well plate filled with PBS, and treated with 5% normal donkey serum (NDS) and 0.3% Triton X-100 in PBS for 60 min. Sections were incubated overnight at 4°C in 5% NDS and 0.3% Triton X-100 in PBS with antibodies to parvalbumin (anti-PV in mouse, 1:4000, Swant Cat# 235, RRID:AB_10000343), calbindin (anti-CB in rabbit, 1:250, Swant Cat# CB38, RRID:AB_2721225), calretinin (anti-CR in goat, 1:4000, Swant Cat# CG1, RRID:AB_10000342), reelin (anti-reelin in mouse, 1:1000, Millipore Cat# MAB5364, RRID:AB_2179313), neuropeptide Y (anti-NPY in rabbit, 1:1000, ImmunoStar Cat# 22940, RRID:AB_2307354), nitric oxide synthase (anti-nNOS in rabbit, 1:500, Millipore Cat# AB5380, RRID:AB_91824), or somatostatin (anti-SOM in rat, 1:1000, Millipore Cat# MAB354, RRID:AB_2255365). After three washing steps in 1% NDS and 0.2% Triton X-100 in PBS, sections were incubated with a FITC-conjugated secondary antibody (anti-rabbit or anti-goat in donkey, or anti-rat or anti-mouse in goat, as appropriate, 1:200; Molecular Probes Cat# A-21206, RRID:AB_141708, Cat# A-11055, RRID:AB_142672, Cat# A-11029, RRID:AB_138404, Cat# A-11006, RRID:AB_141373) in 5% NDS and 0.3% Triton X-100 in PBS for 2 h. Cell nuclei were labeled with Hoechst (1:5.000, Sigma-Aldrich) in 1% NDS and 0.2% Triton X-100 in PBS for 15 minutes. Sections were rinsed twice, mounted with FluorSave and sealed with coverslip. A confocal microscope was used to capture z-stacks of images in regions of interest in CA1 containing the retrograde tracer and FITC signal (Olympus Fluoview FV10i, 10× objective lens, N/A 0.4, 1024 x 1024 pixel, z-step size of 2 or 5 μm). Slices were tested for one antibody. Double-labeling of tracer and antibody staining was analyzed at the focal plane of retrogradely labeled neurons.

For amplification of hM4D(Gi)-mCherry signals, sections containing the RSC and the injection site (i.e., CA1 or AV) were prepared using the same method as above, and were washed in 1% H_2_O_2_ (v/v in methanol) and 5% goat serum (v/v in PBS) before being incubated overnight in chicken anti-mCherry antibody (1:16000 v/v in PBS, Abcam Cat# ab205402, RRID:AB_2722769). Sections were washed in 1% normal goat serum (v/v in PBS) and incubated in biotinylated anti-chicken IgG (H+L) made in goat (1:200 v/v in PBS, Vector Laboratories Cat# BA-9010, RRID:AB_2336114) for 1 hour. The ABC kit (Vector Laboratories Cat# PK-6100, RRID:AB_2336819) was used to amplify the immunosignal. Sections were further incubated in 3,3’-Diaminobenzidine (DAB; 1 tablet in 5 mL H_2_O, Sigma, D4293-50SET) for ~7 mins, before being mounted on a slide and sealed with FluorSave and a #1 coverslip.

The DAB staining was visualized using a macroscope (Olympus SZX16, equipped with a QImaging Retiga2000R camera). Images were acquired using Ephus software, without spatial binning.

***In vivo* photostimulation and electrophysiology.** The *in vivo* methods were recently described in detail ^65^. Briefly, mice first underwent injection of AAV carrying ChR2 into either CA1 or AV, using the same coordinates and methods as described above for *ex vivo* experiments. At least three weeks later, isoflurane-anesthetized mice underwent placement of a head-post over the posterior fossa. A small ( 1 mm) craniotomy was opened over the right RSC. Anesthesia was transitioned to ketamine-xylazine (initial dose: ketamine 80–100 mg/kg, xylazine 5–15 mg/kg, intraperitoneal). Mice were transferred to the recording rig, head-fixed, and thermally supported with heating pad (DC Temperature Control System, FHC). Anesthesia level was continually monitored based on whisking activity and reflexes (toe-pinch), and additional doses of ketamine/xylazine were given subcutaneously as needed to maintain a stable plane of anesthesia. ACSF was applied to keep exposed brain areas moist. An LED-coupled optical fiber (Thorlabs, FG400AEA  400 µm core multimode fiber, 0.22 NA, 6.1 mW output measured at the tip), mounted on a motorized micromanipulator, was positioned over the RSC, and photostimuli (10 ms-duration light pulses) were delivered every 2 s. Trials were repeated 30 or more times. A two-shank linear array (coated with DiI, 0.5 mm separation between shanks, each with 16 channels with 1-2 MΩ impedance and 50-μm spacing; model A2 × 16–10mm-50–500-177-A32, NeuroNexus, Ann Arbor, MI), mounted on a motorized 4-axis micromanipulator, was positioned at the cortical surface (coordinates of the anterior-most shank: -1.6 mm caudal to bregma and 0.3-0.4 mm lateral to midline), and slowly inserted into the cortex to a depth of 1 mm from the pia. Signals were amplified using an RHD2132 digital electrophysiology amplifier board (Intan). We used a RHD2000 USB Interface Evaluation Board (Intan) streaming in the linear array data from the amplifier and piping the data stream in or out of the PC via USB cable. Recorded data, sampled at 30 KHz, were stored as raw signals from the amplifiers and filtered by a 60 Hz notch filter. A digital high-pass filter (800 Hz cutoff, second-order Butterworth) was used to shrink the photovoltaic artifact to the first 3ms post-stimulus window. We analyzed multi-rather than single-unit activity; single units could not be reliably isolated in within the barrages of activity evoked by optogenetic stimulation ^65^. For this, a threshold detector was applied with threshold set to the 4 times the standard deviation to detect spike-like events. To mask the photovoltaic effect, spike counts of the 3 ms window from stimulus onset and offset were then replaced by null values (not-a-number, NaN, in Matlab). These data from each shank were then combined and used to construct peristimulus time histograms, averaged across trials ^65^. After each recording, mice were immediately sacrificed for post-hoc analysis of recording position and injection site. Mice with misplacement in any of these were excluded from analysis.

**Contextual fear conditioning and chemogenetic silencing.** Thirty-eight mice, including 17 for vehicle (ACSF) and 21 for CNO infusion, were used for thalamic axonal silencing experiments. Twenty-nine mice, including 14 for vehicle and 15 for CNO infusion, were used for CA1 axonal silencing experiments. One week before the start of fear conditioning, mice previously injected with hM4D(Gi)-mCherry into either CA1 or thalamus were anesthetized with 1.2% tribromoethanol (v/v, Avertin) ^13^ and craniotomy was performed bilaterally above the RSCs. A guiding cannula (bilateral, short pedestal (5 mm), 26 gauge, 0.8 center-to-center distance, cut 0.75 mm below pedestal; PlasticOne) was slowly inserted into RSC and affixed to cranium using carboxylate cement (Durelon, 3M ESPE). A dummy cannula was inserted and a screw cap was attached to the pedestal to prevent clogging. Mice were single housed in cage and kept in a ventilated cabinet (Scantainer, Scanbur) for 1 week before the day of experiment.

For contextual fear conditioning, mice were briefly exposed, for 3 min, to context A: a 35 × 20 × 20 cm black box with a grid floor, an ethanol odor, and the room light on. This exposure was immediately followed by a foot shock (2 s, 0.7 mA, constant current). This procedure results 40-60% freezing, allowing to observe modulatory effects of chemogenetic manipulations in either direction. Fourty-eight hours later, mice were tested for fear memory retrieval to context, by placing them in context A for 3 mins. Freezing was scored every 10 s by different observers unaware of the experimental condition during context exposures, and expressed as a percentage of the total number of observations during which the mice were motionless with crouching posture. All behavioral experiments were performed between 10 am and 2 pm. Littermates were randomly assigned to the different treatment conditions (CNO or vehicle infusion, see below). All behavioral tests were performed with experimenters blind to genotypes and drug treatments.

Clozapine-N-oxide (CNO, Sigma; 1 mM) or vehicle was infused into RSC 30 mins before fear conditioning. The efficacy of this CNO concentration for silencing synaptic transmission at terminals was previously shown for subcortical ^66^ and RSC circuits ^13^ (and see below for additional characterizations). Mice were anesthetized with isoflurane, and the dummy cannula was replaced with an internal cannula connected to Hamilton microsyringes. An automatic microsyringe pump controller (Micro4-WPI) was used to infuse CNO or ACSF at a rate of 0.5 μL/min until a volume of 0.20 μL per side was reached. In control experiments, when fluorescent muscimol was infused at this rate, the fluorescent signal was confined to the RSC, spanning layers 1 through 6 (**Fig. S4J**). After infusion, mice were placed back to home cage for recovery before conditioning.

Post-hoc histological analysis was performed to confirm accurate targeting of the AAV to dorsal CA1 or AV (**Fig. S4K,L**). After the test, mice were perfused and their brains were fixed and sectioned for mCherry amplification (see Immunohistochemistry section above). Sections were examined for cannula placement, injection site, and axonal projection pattern. Mice with a misplaced cannula, mis-targeted injection site, or weak mCherry expression were excluded from analysis (n = 6 mice for CA1 silencing, n = 11 mice for ATN silencing).

To control for the specificity of CNO and assess its efficacy in reducing synaptic transmission of hM4D(Gi) expressing CA1-RP→RSC and ATN-TC→RSC pathways, we injected AAV virus encoding hChR2 with or without AAV virus encoding hM4D(Gi) into CA1 of Gad2-Cre mice (Cre-dependent virus, 2:5 v/v ratio for hChR2 and hM4D(Gi)) or into thalamus of WT mice (Cre-independent virus, 1:5 v/v ratio). In RSC slices, wide-field photostimulation with an LED was used to activate ChR2-expressing axons while recording evoked synaptic currents from L5pyr neurons every 30 s. After recording a stable baseline for 5 mins, CNO was added at an initial concentration of 0.1 μM, previously shown to be effective in hyperpolarizing hM4D(Gi)-expressing somata ^49^ or silence synaptic transmission in other pathways ^13, 51^. After 7.5 mins, the CNO concentration was increased to 1 μM ^51^. We have previously shown that CNO infusion into the RSC of wild-type mice (i.e., without hM4D(Gi) expression) did not induce off-target effects during CFC ^13^.

**Drugs.** SR-95531 hydrobromide (Gabazine, #1262), CGP55854 hydrochloride (#1248), QX314 chloride (#2313), NBQX (#0373), (*R*)-CPP (#0247) and TTX (#1069) were purchased from R&D Systems. 4-AP (275875-1G) and CNO (C0832-5MG) were purchased from Sigma-Aldrich. For *ex vivo* experiments, all drugs were dissolved in H_2_O, except NBQX which was dissolved in DMSO. For *in vivo* experiments, CNO was dissolved in ACSF.

**Statistical analysis.** Statistical analysis was performed using standard Matlab functions. Non-parametric tests such as signed-rank (for paired data) or rank sum (unpaired data) tests were used to compare groups, unless otherwise noted, with significance defined as p < 0.05. The mean ± s.e.m., unless otherwise noted, were used as statistical measures of central tendency and dispersion.

**Competing interests:**

The authors declare no competing interests.

**Author Contributions:**

N.Y., J.R., and G.S. designed research; N.Y., X.L., and L.L. performed electrophysiology, imaging, immunostaining, and reconstruction; L.R. and J.R. performed behavioral studies; and N.Y., J.R, and G.S. wrote the paper.

## ACKNOWLEDGEMENTS

We thank N. Bernstein, A. Guedea, and D. Wokosin for a technical assistance. We thank A. Apicella, J. Barrett, K. Guo, K. Harris, G. Maccaferri, and A. Tanimura for comments and suggestions. This research was supported by grants from the National Institutes of Health, including the National Institute of Neurological Disorders and Stroke (NS061963), the National Institute of Mental Health (MH108837), and the National Institute of Biomedical Imaging and Bioengineering (EB017695).

## SUPPLEMENTAL INFORMATION

**Figure S1.**
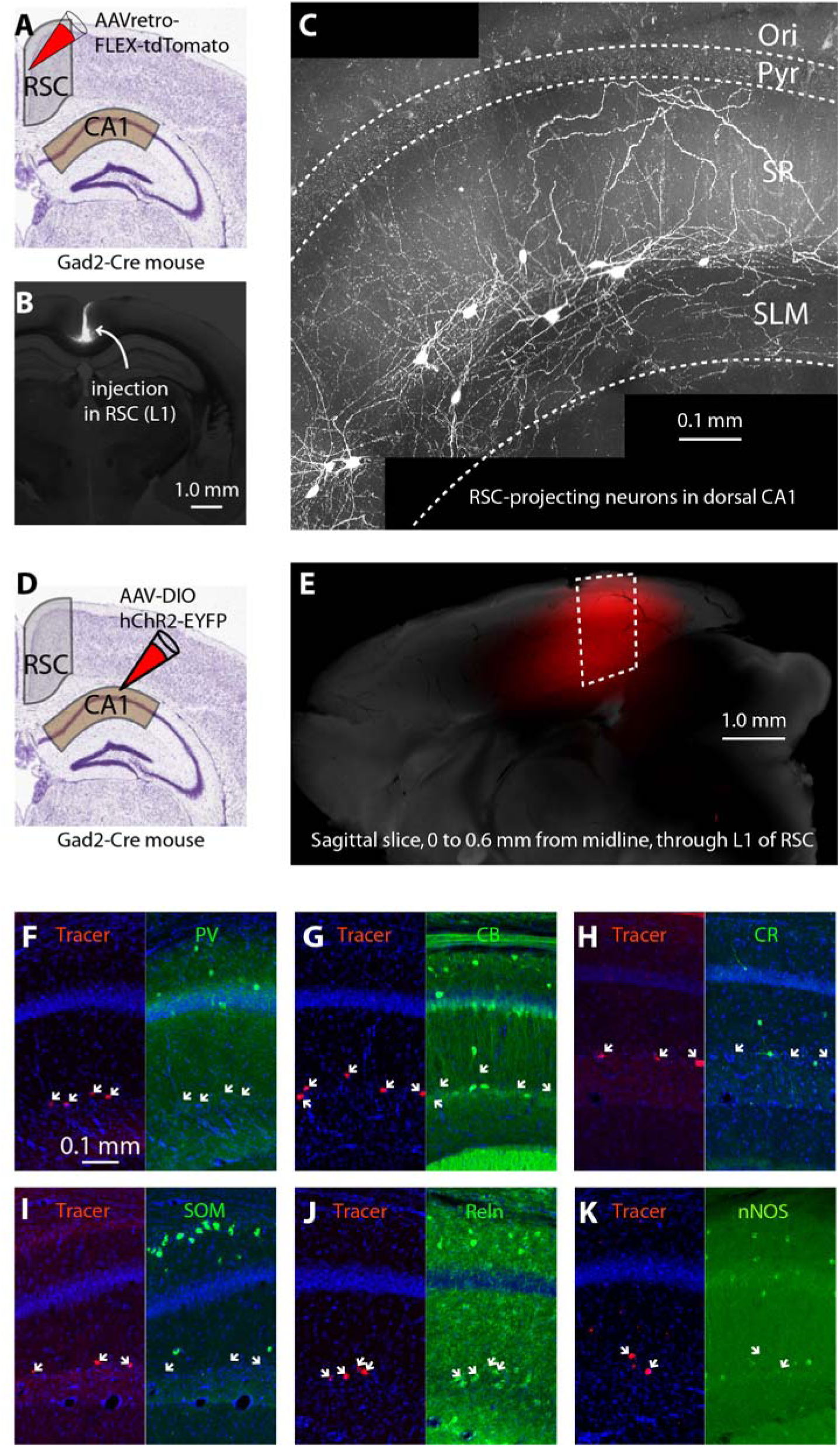
Additional characterizations of CA1-RP neurons and axons. **(A)** Schematic depicting injection of AAVretro-FLEX-tdTomato into L1 of the RSC in a Gad2-Cre mouse, for retrograde labeling of CA1-RP neurons. **(B)** Merged epifluorescence and bright field image showing the injection site in RSC. **(C)** 2-photon maximum-intensity projection image of dorsal CA1, showing several retrogradely transfected neurons labeled with tdTomato around the SR-SLM border. **(D)** For anterograde labeling of GABAergic projections, AAV5-Ef1a-DIO-hChR2-EYFP was injected into dorsal CA1. **(E)** Merged image of a 0.6 mm-thick sagittal slice immediately adjacent to the midline, through L1 of RSC, showing the antero-posterior extent of the CA1 axons projection (red), which is focused onto the RSC. Brain slice recordings in this study were made from RSC neurons located in coronal sections taken through the region indicated (white dashed box). **(F)** Anti-parvalbumin (PV) immunolabeling. There were 0 immunopositive neurons out of 47 retrogradely labeled neurons (6 dorsal hippocampal fields of view, 2 mice). The scale bar also applies to other panels. **(G)** Anti-calbindin (CB) immunolabeling. There were 0 immunopositive neurons out of 32 retrogradely labeled neurons (3 dorsal hippocampal fields of view, 2 mice). **(H)** Anti-calretinin (CR) immunolabeling. There was 1 immunopositive neuron out of 24 retrogradely labeled neurons (4 dorsal hippocampal fields of view, 2 mice). **(I)** Anti-somatostatin (SOM) immunolabeling. There were 0 immunopositive neurons out of 29 retrogradely labeled neurons (5 dorsal hippocampal fields of view, 2 mice). **(J)** Anti-reelin (Reln) immunolabeling (see main text for details). **(K)** Anti-NOS immunolabeling (see main text for details).

**Figure S2.**
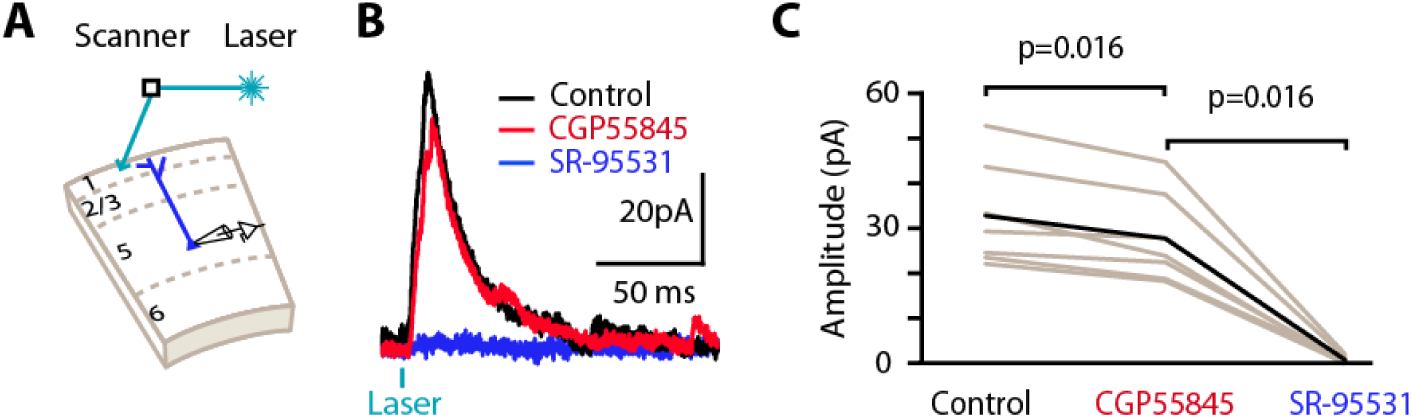
Involvement of GABA_B_ receptors in CA1-RP inhibitory synaptic transmission to L5pyr neurons. **(A)** Laser scanning photostimulation was used to activate CA1-RP axons in L1 of RSC. **(B)** Example traces of photo-evoked IPSCs after sequential application of GABA_B_ receptor antagonist CGP55845 (red, 5 μM) and GABA_A_ receptor antagonist SR-95531 (blue, 10 μM). **(C)** Group comparison of drug effect. Gray line indicates data from individual cells and black line indicates mean. CGP55845 reduced IPSC mean amplitude by 16% (control 32.7 ± 4.4 pA, CGP55845 27.6 ± 3.8 pA, signed-rank test, n = 7 cells). The remaining responses were eliminated after addition of SR-95531 (0.5 ± 0.4 pA).

**Figure S3.**
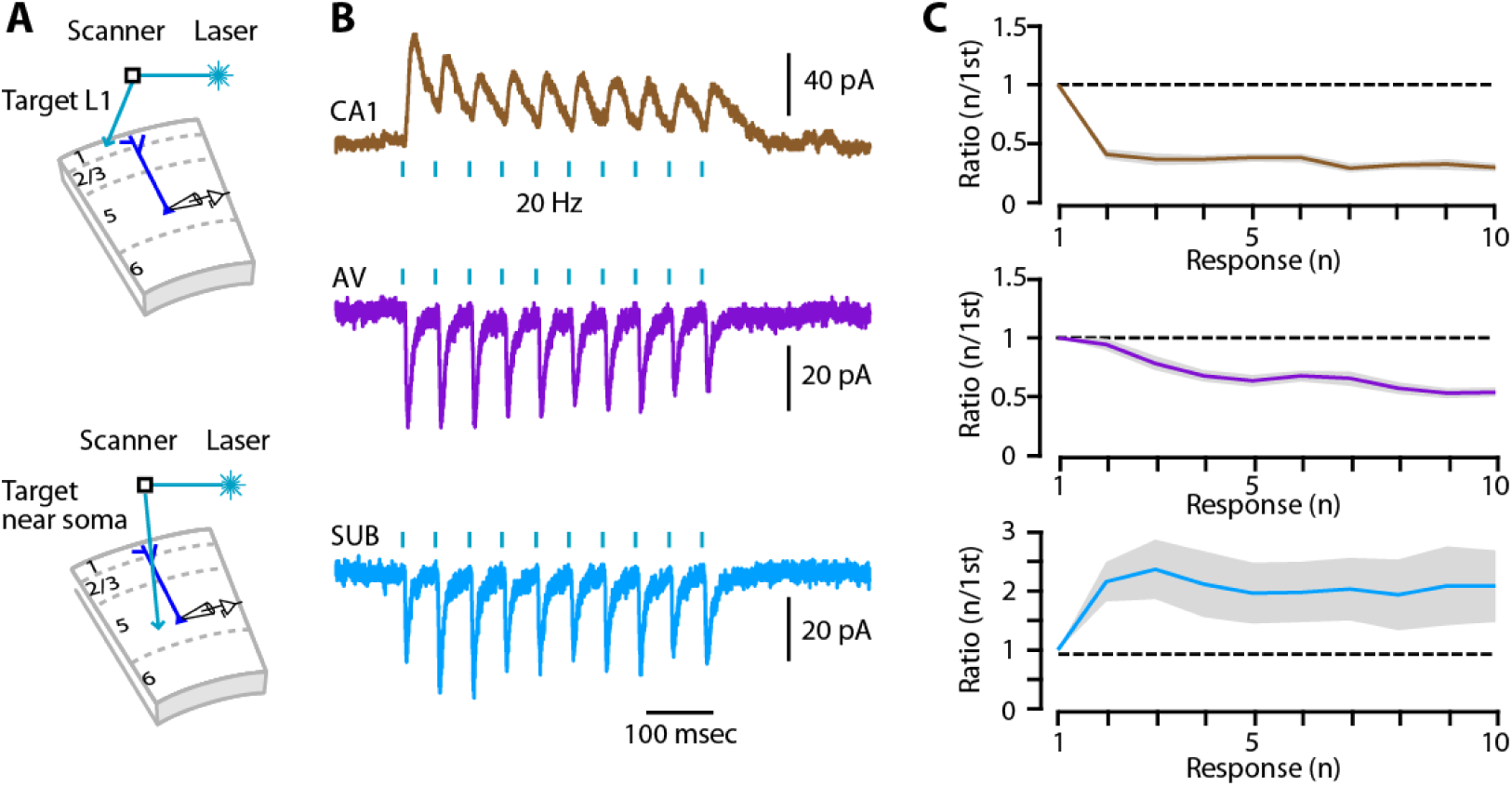
Short-term synaptic plasticity of CA1, ATN, and subicular synapses terminating on L5pyr neurons. **(A)** Schematics showing approach used to repetitively stimulate CA1 and ATN axons (laser aimed at sites in L1; top) or subicular (SUB) axons (laser aimed at sites in deeper layers; bottom). **(B)** Example traces showing responses of recorded in L5pyr neurons to 20 Hz repetitive stimulation of CA1, ATN and SUB inputs. (**C**) Group data. Top, analysis of CA1 inputs, showing pattern of strong synaptic depression (p = 0.006 or less for all responses when compared against 1^st^ response, signed-rank test, Bonferroni-corrected, n = 26 L5pyr neurons). Middle, analysis of ATN inputs, showing initially non-depressing followed by depressing responses (p = 0.006 or less for 3^rd^ to 10^th^ response when compared against 1^st^ response, signed-rank test, Bonferroni-corrected, n = 18 L5pyr neurons). Bottom, analysis of subicular inputs, showing facilitating responses (p = 0.006 or less for 2^nd^ and 3^rd^ response vs 1^st^ response, signed-rank test, Bonferroni-corrected, n = 23 L5pyr neurons).

**Figure S4.**
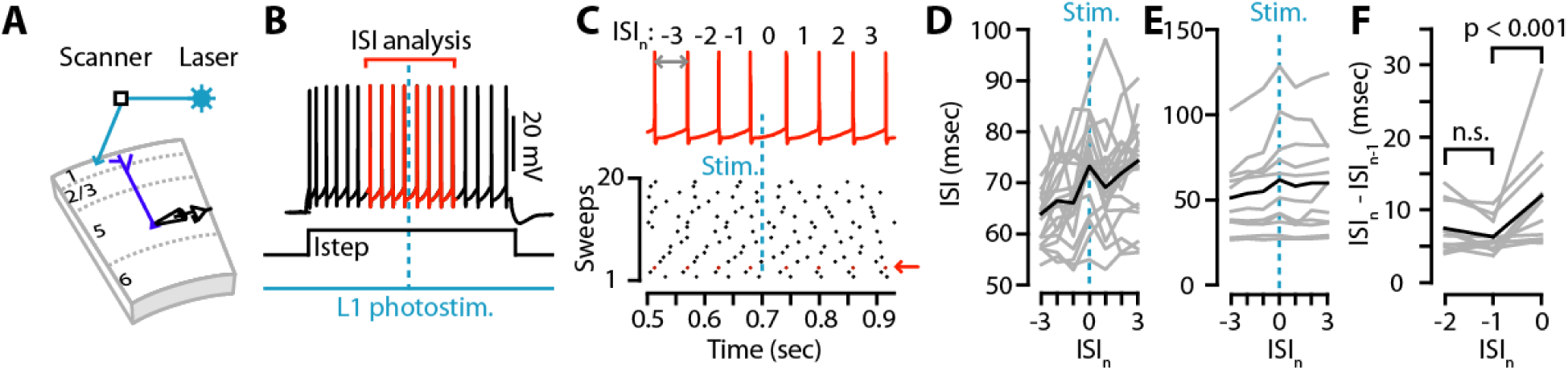
Effect of CA1 inhibitory input on repetitive firing of L5pyr neurons. (**A**) Schematic of the experimental strategy, involving whole-cell recording from an RSC-L5pyr neuron while briefly flashing the focused beam of the laser to activate CA1 axons at a location slightly away from the dendrites of the recorded cell. (**B**) Example trace, showing a train of spikes evoked in a L5pyr neuron by a step of current injected via the somatic patch pipette (1 s, +200 pA), and the timing of the laser pulse (1-ms duration) used to photostimulate CA1 axons. Subsequent analysis of inter-spike intervals (ISI) focused on the peristimulus region indicated by red. (**C**) Top: Enlarged peristimulus spiking shown in panel B. Bottom: The spiking patterns recorded in multiple sweeps are shown bottom; the red arrow indicates the sweep of the example trace. (**D**) Plot of peristimulus ISIs for one neuron, showing the mean (black) and individual sweeps (gray). The zeroeth ISI is defined as the one during which the laser stimulus was delivered. (**E**) Plot of the mean peristimulus ISIs for multiple neurons (grey), and the overall mean across all neurons (black). (**F**) Plot of the change in ISI, calculated by subtracting the preceding ISI, for multiple (n = 11) neurons (grey) and the overall mean across all neurons (black). The mean change in ISI did not differ significantly for the two ISIs immediately before the L1 stimulation (2.4 ± 1.0 ms vs 1.3 ± .7 ms, p = 0.12, signed-rank test), but did for the ISI before vs during stimulation (1.3 ± 0.7 ms vs 6.9 ± 2.2 ms, p = 0.003, signed-rank test).

**Figure S5.**
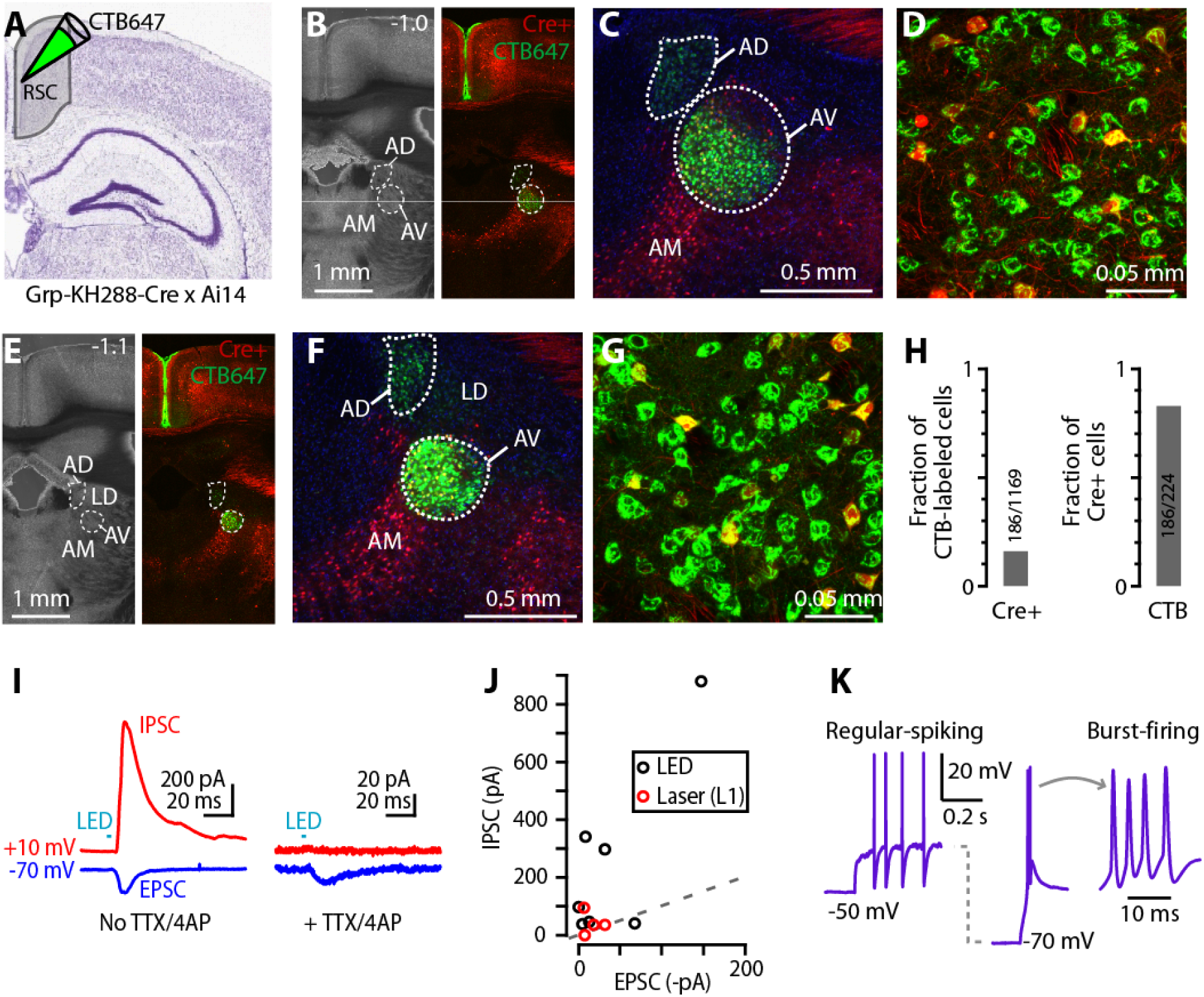
Additional characterizations of ATN→RSC circuits. **(A)** Characterization of thalamic labeling in Grp_KH288-Cre mice. CTB647 was injected into L1 RSC of Grp_KH288-Cre mice crossed with a fluorescent reporter line (Ai14). **(B)** Bright-field (left) and epifluorescence (right) image of a coronal brain slice (1.0 mm posterior to bregma) containing anterior thalamic nuclei, including AV and AD (dashed circle). Retrograde labeling is also shown (green). **(C)** Higher-magnification view. Cre-positive neurons are present in AV, but not in AD. (**D**) Higher-magnification view of the labeling in AV in (C). A subset of RSC-projecting neurons is Cre-positive. **(E)** Coronal brain slice (1.1 mm posterior to bregma, immediately posterior to that shown in B), containing anterior thalamic nuclei, including AV and AD (dashed circle). **(F)** Higher-magnification view. Retrograde labeling is also shown (green). **(G)** Higher-magnification view of the labeling in AV. A subset of RSC-projecting neurons is Cre-positive. (**H**) Left: Quantification of double-labeled neurons relative to the CTB-labeled neurons in AV (analyzed 4 slices from 2 mice, 2 slices per mice). Right: Quantification of double-labeled neurons relative to the Cre+ neurons. **(I)** Thalamocortical excitation and feedforward (disynaptic) inhibition. Example traces from a L5pyr neuron in an RSC brain slice, showing responses to photostimulation of ATN-TC axons while the voltage was held at either -70 mV (blue traces) to sample inward currents (EPSCs) or at +10 mV (red traces), to sample outward currents (IPSCs). Addition of TTX and 4-AP eliminated the IPSC but not the EPSC, indicating that the IPSC is mediated by feedforward (disynaptic) mechanism. **(J)** Comparison of EPSC and IPSC mean amplitude. ATN-TC axons were stimulated by wide-field LED (black circle) or focal stimulation of L1 using a laser. (**K**) Example traces, recorded from a RSC-projecting TC neuron in AV, showing regular-spiking vs bursting firing patterns evoked by somatic injection of a current step stimulus from depolarized (-50 mV) versus hyperpolarized (-70 mV) membrane potentials. Trace on the right shows the middle trace’s burst on a faster time base.

**Figure S6.**
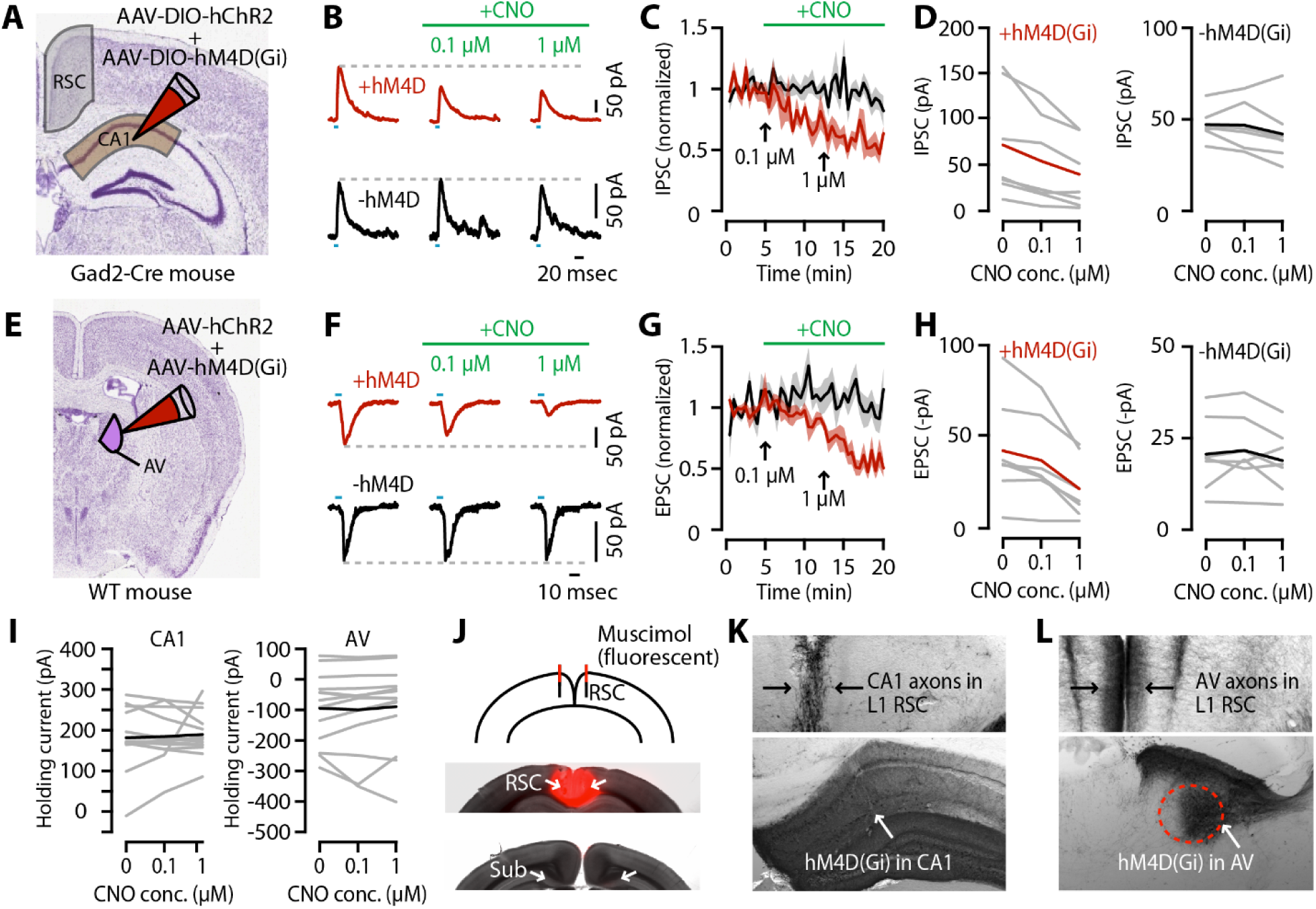
Chemogenetic control experiments. **(A)** Schematic of the injection strategy for assessing effect of CNO on synaptic transmission from presynaptic hM4D(Gi)-expressing CA1-RP axons to postsynaptic RSC pyramidal neurons. AAV5-Ef1a-DIO-hChR2-EYFP and AAV8-hSyn-DIO-hM4D(Gi)-mCherry were co-injected into CA1 of Gad2-Cre mice. **(B)** Example traces showing photo-evoked responses at baseline (left) and increasing concentrations of CNO (right), for axons that either did (top row, +hM4D) or did not (bottom row, -hM4D) co-express the chemogenetic construct. **(C)** Time course of the amplitude (mean ± s.e.m., normalized to baseline) of the optogenetically evoked response, for axons that either did (red, n = 7) or did not (black, n = 7) co-express the chemogenetic construct. **(D)** Group comparison of photo-evoked IPSC amplitudes recorded with different concentrations of CNO. Left: Synaptic transmission from terminals with hM4D(Gi). The mean IPSC amplitude 2.5 mins immediately before application of 0.1 μM CNO was 70.7 ± 21.9 pA. The mean IPSC amplitudes 2.5 mins before application of 1 μM CNO and 2.5 mins before the end of recording session were 53.4 ± 17.7 pA and 39.2 ± 13.6 pA, respectively (baseline vs 0.1 μM or1 μM CNO: n = 7, p = 0.016 for both, signed-rank test; 0.1 μM CNO vs 1 μM CNO, p = 0.03, signed-rank test). Right: Synaptic transmission from terminals without hM4D(Gi), measured at same time window as plot on the left. The mean IPSC amplitude at 0, 0.1 and 1 uM CNO was 47.2 ± 3.2, 46.8 ± 4.7, and 42.1 ± 6.0 pA, respectively (baseline vs 0.1 μM or1 μM CNO: n = 7, p = 0.94 or 0.16, signed-rank test; 0.1 μM CNO vs 1 μM CNO, p = 0.11, signed-rank test). **(E-G)** Same, for assessing effect of CNO on synaptic transmission from presynaptic hM4D(Gi)-expressing ATN-TC axons to postsynaptic RSC pyramidal neurons. Cre-independent AAV1-CamKIIa-hChR2-mCherry and AAV8-hSyn-HA-hM4D(Gi)-mCherry were injected into AV of WT mice. (**H**) Left: Synaptic transmission from terminals with hM4D(Gi). The mean EPSC amplitude 2.5 mins immediately before application of 0.1 μM CNO was 42.1 ± 10.7 pA. The mean EPSC amplitudes 2.5 mins before application of 1 μM CNO and 2.5 mins before the end of recording session were -36.9 ± 9.2 pA and 21.5 ± 6.3 pA, respectively (baseline vs 0.1 μM or 1 μM CNO: n = 7, p = 0.031 or 0.016, signed-rank test; 0.1 μM CNO vs 1 μM CNO, p = 0.031, signed-rank test). Right: Synaptic transmission from terminals without hM4D(Gi). The mean EPSC amplitude at 0, 0.1 and 1 uM CNO was -20.5 ± 3.8, -21.5 ± 3.7, and -18.8 ± 3.2 pA, respectively (baseline vs 0.1 μM or1 μM CNO: n = 7, p = 0.81 or 0.16, signed-rank test; 0.1 μM CNO vs 1 μM CNO, p = 0.16, signed-rank test). **(I)** Group comparison of holding current recorded with different concentrations of CNO. Left: Mean holding current during recording of CA1 input in absence of CNO was 180.3 ± 20.1 pA. The mean holding current during recording with 0.1 and 1 μM CNO was 183.0 ± 16.1 and189.4 ± 14.6 pA (baseline vs 0.1 μM or1 μM CNO: n = 14, p = 0.855 and 0.808, signed-rank test; 0.1 μM CNO vs 1 μM CNO, p = 0.626, signed-rank test). Right: Mean holding current during recording of ATN input in absence of CNO was -96.0 ± 30.4 pA. The mean holding current during recording with 0.1 and 1 μM CNO was -100.9 ± 35.5 and -89.7 ± 35.9 pA (baseline vs 0.1 μM or1 μM CNO: n = 14, p = 0.903 and 0.153, signed-rank test; 0.1 μM CNO vs 1 μM CNO, p = 0.104, signed-rank test). **(J)** Merged bright-field and epifluorescence image showing restriction to the RSC of a fluorescent dye (fluorescent muscimol) that was cannula-infused into the RSC using the same parameters used for the behavioral experiments. No labeling of dye was observed in dorsal subiculum. **(K)** mCherry expression at the injection site in CA1, and in RSC (axonal labeling). Arrow indicates region containing labeled cells. **(L)** mCherry expression at the injection site in AV, and in RSC (axonal labeling). Arrow indicates region containing labeled cells.

**Table S1.** Intrinsic properties of CA1-RP neurons, related to Fig. 1. R_in_, input resistance; V_r_, resting membrane potential; AP, action potential; f/I, frequency/current.

|  | **R<sub>in</sub>**<br>**(MΩ)** | **V<sub>r</sub>**<br>**(mV)** | **AP**<br>**threshold**<br>**(mV)** | **AP**<br>**height**<br>**(mV)** | **AP**<br>**width**<br>**(ms)** | **Rheobase**<br>**(pA)** | **Sag**<br>**(%)** | **f/I slope**<br>**(Hz/nA)** | **Max. spike**<br>**frequency**<br>**(Hz)** |
| --- | --- | --- | --- | --- | --- | --- | --- | --- | --- |
| **N** | 24 | 24 | 24 | 24 | 24 | 10 | 14 | 24 | 14 |
| **Mean** | 246.4 | -63.6 | 36.7 | 57.5 | 0.75 | 74.0 | 16.6 | 204.2 | 54.7 |
| **S.D.** | 78.1 | 5.2 | 5.2 | 7.6 | 0.20 | 26.7 | 6.6 | 59.8 | 18.9 |

